# Early-life conditions shape baseline immune gene expression, but not pathogen-induced immune activation, in brown trout

**DOI:** 10.64898/2026.09.11.750927

**Authors:** Quinn Alexander Coxon, Stephanie C. Talker, Helena Saura Martinez, James Ord, Nicolas Diserens, Heike Schmidt-Posthaus, Irene Adrian-Kalchhauser

## Abstract

Early-life environment and parental background can shape immune phenotypes, but whether such differences persist during infection and influence the immune response remains unclear. We used single-cell RNA sequencing to characterize kidney immune responses in brown trout (*Salmo trutta*) from three origins differing in parental history and early rearing: wild parents with offspring reared in the wild; wild parents with offspring reared in a hatchery; and hatchery parents with offspring reared in a hatchery. All fish originated from the same river and showed no detectable genetic population structure. Fish were exposed to *Tetracapsuloides bryosalmonae*, the causative agent of proliferative kidney disease, and analyzed 25 days later alongside unexposed controls.

Across 18 scRNA-seq datasets, we identified 30 cell clusters and found a clear transcriptional imprint of developmental and parental history in the resting immune system. In unexposed fish, origin was associated with pronounced differences in gene expression, particularly in B cells. When fish were subsequently exposed to the pathogen and developed subclinical infections, both immune-cell composition and transcriptional state experienced pronounced changes: T cells increased in relative abundance, while B cells, neutrophils and proliferating progenitors showed the strongest transcriptional responses, consistent with an active but controlled subclinical response. Strikingly, the strong origin-dependent differences present before exposure largely disappeared after infection. Fish from all three backgrounds converged on a shared transcriptional response, and genes associated with origin showed little overlap with those responding to infection. Thus, at single-cell resolution in vivo, we identify infection-induced transcriptional convergence across distinct environmental and parental backgrounds. Our findings show that substantial baseline immune variation can persist under resting conditions yet be overridden by pathogen exposure, constraining distinct immune states towards a common response.

## Introduction

Environmental challenges experienced during development and living conditions of the parental generation contribute to phenotypic responses of individuals and can alter reaction norms, including the magnitude, timing, and cellular composition of responses to pathogens ^1, 2^. Understanding these impacts of early-life and parental experiences on immune responses is important in conservation management because individuals released from captivity may respond differently from wild individuals when exposed to disease ^3^, matters to livestock production for similar reasons, and has implications for evolutionary biology because such environmentally induced variation may influence fitness differences, selection and adaptation across prolonged timescales ^4, 5^.

Teleost fishes provide a particularly relevant system in which to investigate these effects. Wild populations face increasing disease pressure associated with habitat degradation, pollution, climate change, stocking, trade, and species introductions, all of which can promote pathogen emergence and transmission ^6, 7^. In aquaculture, infectious diseases are a major constraint on food security, animal welfare, and sustainable production ^8^. Understanding how developmental history and parental background shape immune responses is therefore directly relevant to both the conservation of wild fish populations and the management of cultured stocks.

Brown trout (*Salmo trutta*) provide a particularly relevant system in which to examine these effects because early-life and parental influences intersect directly with current management practices. Populations are frequently supplemented with hatchery-reared fish, including in rivers affected by disease. In Switzerland, more than seven million hatchery-reared trout are released annually ^9^, yet stocked fish often make only a limited contribution to adult populations (^10–12^, own unpublished observations), suggesting that hatchery-associated conditions may alter traits important for survival after release. Indeed, captive breeding and early rearing can affect physiology and gene regulation, including immune function ^13–15^. while parental background may introduce additional variation. Consistent with this, we previously identified baseline gene-expression differences among immune cells from wild- and hatchery-associated brown trout using scRNA-seq ^16^, as well as an origin-specific increase in T-cell abundance following *T. bryosalmonae* exposure using flow cytometry ^17^. What remains unclear is whether early rearing environment and parental background also matter beyond baseline status and modify the cellular and transcriptional response to infection.

Brown trout are often exposed to *Tetracapsuloides bryosalmonae*, the causative agent of the well-studied Proliferative Kidney Disease (PKD) ^18, 19^. Infection outcomes vary depending on affected fish species and environmental context. Under elevated temperatures >15°C or additional stress, like pollution, PKD can cause severe renal hyperplasia and granulomatous nephritis, anemia, and mortality ^20–24^. Under more favorable conditions, infected fish may lack overt clinical signs despite persistent parasite detection and substantial immunological activity ^17, 25^. Young-of-the-year (YOY) salmonids are particularly susceptible during their first exposure, whereas older fish often develop partial acquired protection ^26^. Gene-expression analyses suggest that severe disease is associated with an excessive or poorly controlled immune response, including pronounced B-cell proliferation and immunoglobulin production ^25, 27, 28^, whereas mild infection has been associated with limited changes in immune cell composition ^17^. Because disease outcome depends critically on external factors, PKD provides a well-suited system for testing whether developmental and parental background modify the cellular and transcriptional response to infection.

Fish immunology remains constrained by the limited availability of cell-type-specific antibodies and other reagents. Single-cell RNA sequencing (scRNA-seq) provides a powerful alternative by resolving immune-cell populations and functional states directly from their transcriptional profiles. In zebrafish, scRNA-seq has mapped the cellular diversity of the kidney and hematopoietic system ^29^, and identified previously unrecognized innate lymphoid-like populations and their responses to immune stimulation ^30^. In salmonids, it has revealed heterogeneity among peripheral blood B cells in rainbow trout ^31^, while single-nucleus transcriptomics has resolved cell-type-specific hepatic responses of Atlantic salmon to *Aeromonas salmonicida* infection ^32^. In brown trout, scRNA-seq has further revealed transcriptional divergence among duplicated immune genes and baseline expression differences associated with rearing environment and domestication history ^16^.

While scRNA-seq immune system characterizations are increasingly common in non-model organisms, analyses of the immune response that account for developmental and parental background remain uncommon in teleosts. Here, we used scRNA-seq to characterize the kidney immune response of brown trout from three origins differing in parental background and early rearing environment. Fish were either left unexposed or exposed to *T. bryosalmonae* under conditions that produced infection without overt disease. This design allowed us to characterize the cellular response to pathogen exposure independently of extensive tissue pathology and to test whether host background altered this response. We first identified the immune-cell populations and exposure-associated transcriptional states present in the trout kidney. We then determined how *T. bryosalmonae* exposure affected immune-cell composition and cell-type-specific gene expression. Finally, we tested whether parental background and early rearing environment influenced baseline immune states and the response to exposure. Together, our results demonstrate the value of single-cell transcriptomics for investigating immune responses in a non-model vertebrate and clarify how developmental history interacts with pathogen exposure in wild- and hatchery-associated fish.

## Methods

### Experimental Design

The experiment was performed under controlled laboratory conditions. Brown trout came from three different rearing conditions: Wild (W), Wild:Farm (W:F) and Farm (F) (**Figure 1**) ^17^.

**Figure 1.**
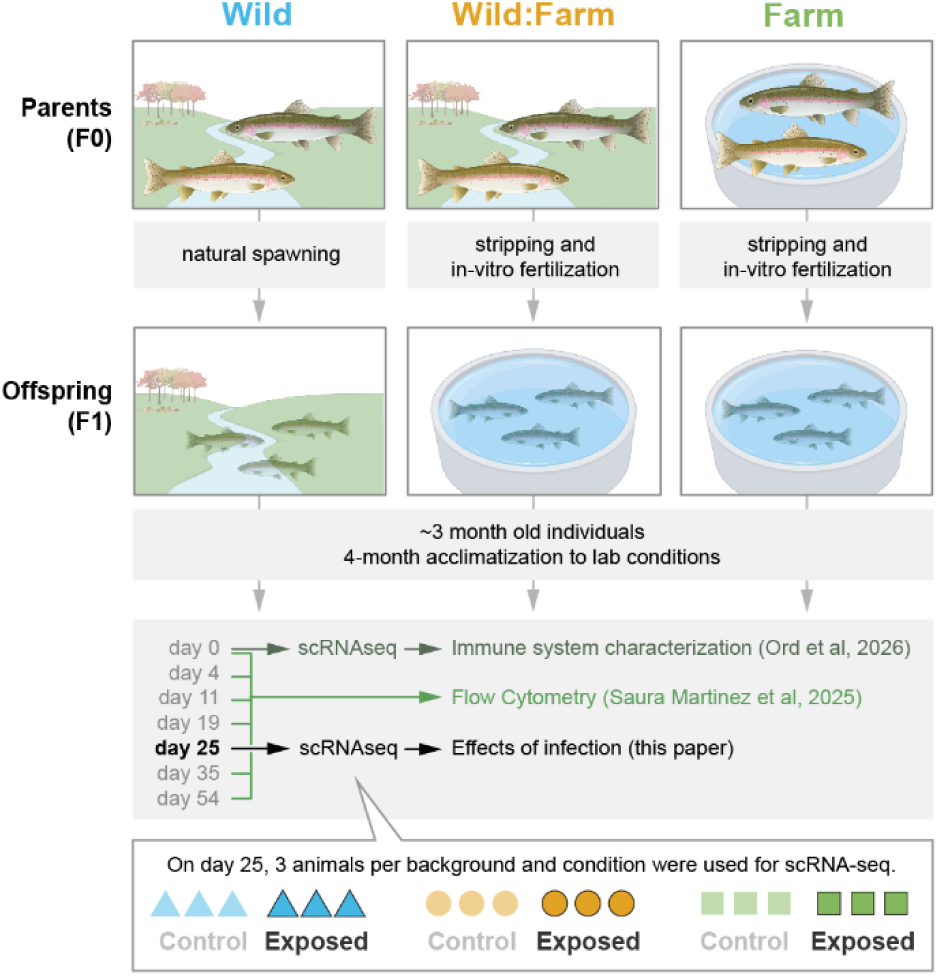
Experimental design and sampling strategy. The immune systems of 7-month old trout from three origins were compared: wild offspring produced by natural spawning and reared in the wild (**Wild**), offspring of wild parents produced by stripping and in vitro fertilization and reared under farm conditions (**Wild:Farm**), and offspring of farm-reared parents produced by stripping and in vitro fertilization and reared under farm conditions (**Farm**). At three months of age, fish were transferred to the laboratory and acclimatized for four months before exposure. Immune-system characterization was performed by scRNA-seq on day 0 ^16^, flow cytometry at all timepoints ^17^ except day 25, and scRNA-seq on day 25 (subject of this study). On day 25, three non-exposed and three exposed fish from each origin (one individual per experimental batch and condition) were selected for scRNA-seq, resulting in 18 experimental individuals in total.

Briefly, the **Wild** group (W) consisted of YOY brown trout that were electrofished in April 2021 in two Swiss rivers of the same river system known to be positive for PKD (Schmidt-Posthaus, own investigations) over a stretch of 100 m each (*Brübach*, between 47.471195 °N/ 9.136465 °E and 47.473303 °N/ 9.140526 °E; and *Rörlibadbach,* between 47.473019 °N/ 9.136563 °E and 47.478731 °N/ 9.134755 °E). The parents (F0 generation) and the YOY (F1 generation, experimental animals) were born and raised in the wild. Thus, the parents were presumably exposed to *T. bryosalmonae,* while exposition of experimental animals is unknown. The **Wild:Farm** group (W:F) consists of juvenile brown trout raised in an aquaculture facility. The parents (F0 generation) of these animals were born and raised in the wild, were electrofished in the same PKD-positive river system (*Brübach*, coordinates between 47.462306 °N/ 9.122916 °E and 47.481201 °N/ 9.159397 °E) and spawned in autumn 2020 under aquaculture conditions in a known PKD negative facility. In-vitro fertilized eggs (F1 generation, experimental animals) were raised until spring 2021 in the aquaculture facility. Thus, the parents were presumably exposed to *T. bryosalmonae*, while the experimental animals were not exposed. The **Farm group** (F) consists of YOY trout raised in the same aquaculture facility. At least two former generations of these animals were born and raised in this Swiss fish farm, although their ancestors originated from the same river system (*Brübach*). The parents were spawned in autumn 2020 and in-vitro fertilized eggs (F1 generation, experimental animals) were raised until spring 2021 in the aquaculture facility. Thus, neither parents nor offspring were ever exposed to *T. bryosalmonae*.

All experimental F1 animals (250 animals per group) were transferred to the facility of the Institute for Fish and Wildlife Health (FIWI), University of Bern in May 2021, when fish were approximately 3 months old. They were held in 21 independent separate 38 L tanks, with separated water supply, at constant 16°C. The fish were acclimatized to laboratory conditions for four months, and clinical signs and mortalities were assessed daily.

After acclimatization, each group (W, W:F, F) was distributed into six 38 L aquaria, with n=29 fish per aquarium, with three aquaria for biological non-exposed replicates and three aquaria for exposed replicates. On day 0, two fish per tank were euthanized with an overdose of buffered tricaine (150 mg/L tricaine methanesulfonate; Tricaine PHARMAQ®, Pharmaq) and investigated for an initial health check as well as for presence of *T. bryosalmonae* DNA ^17^, resulting in 27 fish per aquarium at initiation of experiment (**Figure 1**) (also see ^17^ for experimental design).

### Exposure

To expose the treatment groups to the pathogen, freshwater bryozoans were collected from Swiss rivers known to be endemic for the parasite (*Alte Aare,* coordinates: 590781/ 217845 and *Furtbach*, coordinates: 2670258/ 1255599) as previously described ^17^. They were transferred to the FIWI, bryozoa were disrupted by manual grinding to release the spores, and the homogenate kept in original river water at room temperature for 1 h until addition to the aquaria. From 2 mL homogenate, DNA was extracted and qPCR was performed as previously described ^17^ to confirm presence and concentration of *T. bryosalmonae* DNA (1 000–10 000 parasite DNA copies/ µL). For exposure, the water flow was stopped, aeration increased, and tank water lowered to 20% (approx. 7.6 L). The parasite homogenate was diluted to obtain equal volumes of 24 mL of the homogenate containing 2x10^4^ copies of parasite DNA that were added to the nine exposure tanks. After 1.5 h, the water flow was restarted. Non-exposed fish were treated identically except for the addition of parasites. This procedure was repeated on three consecutive days with the same dose ^17^.

### Sampling and cell isolation

For the present study, kidney cells were isolated on day 25 post-exposure from three exposed and three non-exposed animals per group (Wild, Wild:Farm, Farm) (Fig. 1). Fish were euthanized by immersion in an overdose of tricaine (150 mg/L tricaine methanesulfonate; Tricaine PHARMAQ®, Pharmaq). The kidneys were then dissected (head and trunk kidneys together), and leukocytes were isolated as previously described ^17, 16^. The kidney tissue was passed through a 105 µm pore size mesh filter with Leibovitz’s (L-15) medium (Thermofischer, Reinach, Switzerland) containing 10% fetal calf serum. The resulting cell suspension was then layered onto a sterile, isotonic Ficoll gradient (Ficoll-paque Plus, GE Healthcare Bio-Sciences AB, Sweden) with a density of 1.077 g/mL and spun at 750 x g for 40 minutes at 4°C to enrich white blood cells. Leukocytes at the Ficoll/medium interphase were aspirated, washed in L-15 medium and centrifuged at 290 x g at 4°C for 10 minutes. Cell suspensions were resuspended in PBS and kept on ice until further processing. Cell counting and viability assessment were performed using an automated cell counter (DeNovix® CellDrop Automated Cell Counter, NC, USA).

### Single-cell RNA-seq (10x Genomics)

Samples were diluted to a concentration ranging from 700 to 1900 cells/µL and were immediately processed at the Next-Generation Sequencing Platform of the University of Bern. Gel beads-in-emulsion (GEM) generation and barcoding, reverse transcription, cDNA amplification and 3’ gene expression library generation steps were all performed according to the Chromium Next GEM Single Cell 3ʹ Reagent Kits v3.1 (Dual Index) User Guide (10x Genomics CG000315, Rev E) with all stipulated 10x Genomics reagents. Generally, 9-11 µL of each cell suspension (1500-1900 cells/µL) and 32-35 µL of nuclease-free water were used for a targeted cell recovery of 10,000 cells. Next, GEMs were subjected to library construction using the Chromium^TM^ Single Cell 3’ Library Kit v3.1 (10x Genomics). In the first step, reverse transcription was performed, generating cDNA tagged with a cell-specific barcode and a unique molecular index per transcript. Fragments were size-selected using SPRIselect magnetic beads (Beckman Coulter, Brea, CA, USA). Illumina sequencing adapters were then ligated to the size-selected fragments and cleaned up using SPRIselect magnetic beads. The quality of the final library was assessed using an Agilent 2100 Bioanalyzer (Agilent technologies, Amstelveen, The Netherlands). The samples were subsequently sequenced using a NextSeq 550 instrument (Illumina, CA, US) with 150PE chemistry. Samples were distributed across sequencing runs to minimize batch effects. Individual and average sample and sequencing specifications are given in Supplementary Table 1. On average, 5347 cells and 375’057’796 reads were obtained per individual, and 57’029 reads were obtained per cell.

### Alignment and count tables

The raw FASTQ files were processed using CellRanger (version 7.2.0, 10 x Genomics). Reads were aligned to the brown trout reference genome (fSalTru1.1, GCF_901001165.1, augmented with additional gene annotations produced from kidney RNAseq data as described by ^16^) using *cellranger count* and filtered feature matrices along with BAM files were produced.

### SNP Calling

Aligned BAM files from 18 samples were processed to infer pairwise genetic relatedness. Single-nucleotide polymorphisms (SNPs) were called jointly across all samples using MonoVar ^33^, producing a multi-sample VCF containing genotype likelihoods and per-sample allele depth information. The resulting VCF was cleaned to ensure compatibility with downstream analyses and converted to PLINK 2.0 format, retaining only biallelic SNPs with canonical nucleotide alleles (A, C, G, and T). Variants with a minor allele frequency below 5% or missing genotype rates exceeding 20% were excluded prior to relatedness estimation. Pairwise relatedness coefficients were subsequently estimated using the KING-robust method implemented in PLINK 2.0, generating kinship coefficients for all 18 samples (Supplementary Table 2). Samples were then clustered using agglomerative hierarchical clustering with average linkage (UPGMA), based on these coefficients.

### Clustering and assignment of cluster identities

Cells were clustered using the Seurat package version 4.3.0 in R version 4.2.3. Following quality control, cells with fewer than 200 or more than 5,000 detected genes or with >10% mitochondrial transcript content were excluded (Supplementary Figure 1). Haemoglobin transcripts identified as contaminants were removed prior to downstream analyses, though clustering was performed separately without excluding haemoglobin genes to identify red blood cell clusters. Gene expression counts were log-normalized, and expression values were scaled across all genes before principal component analysis (PCA) was performed on the variable genes. A shared nearest-neighbour graph was constructed using the first 20 principal components and clusters were identified at a resolution of 0.5. Uniform Manifold Approximation and Projection (UMAP) was subsequently performed using the first 20 principal components for two-dimensional visualization of the resulting cell clusters.

Firstly, we used a list of genes previously compiled and tested by ^16^, which will be referred to as “prior markers”. These consisted of the trout homologues of putative cell lineage markers found in other species. The ENSEMBL IDs of trout homologues were identified using the biomaRt R package version 2.52.0. We then used a list of markers collected by ^16^, using the FindMarkers()function on the clusters they had identified by the previous method. Combining these two approaches allowed for more robust identification of cell types, especially for the Monocyte/macrophage-like cluster M13.

Finally, we ran FindAllMarkers()on all 31 clusters and compiled tables of genes upregulated in these clusters. The zebrafish and human homologues of these genes were queried using the ensemble python API (get version) and putative gene functions were added from the Zebrafish Information Network (ZFIN) and the Human Gene Nomenclature Committee (HGNC). This allowed for speculation as to the function of clusters which could not be identified by prior or established markers.

Red blood cells were identified by repeating the previous clustering steps without the exclusion of haemoglobin (Hb) and using the FindAllMarkers() function on the clusters which were defined by the highest expression of Hb transcripts. These lists of RBC marker genes were retained and used for annotating the RBC clusters after Hb transcripts were removed.

### Differential Cell Number Analysis

Cell counts were analysed in R using RStudio 2025.09.2+418 and R version 4.5.0 (2025-04-11 ucrt) How About a Twenty-Six ^34^ using a negative-binomial generalized linear mixed-effects model implemented in glmmTMB ^35^. Origin, treatment, and their interaction were included as fixed effects. Random intercepts for individual and experimental batch accounted for repeated measurements from the same fish and variation among batches, respectively. To account for differences among cell clusters, the model included a cluster-specific random intercept and random slopes for origin and treatment, allowing both baseline cell abundance and the effects of origin and treatment to vary among clusters. The model used the nbinom2 parameterization with a log link, in which the variance increases quadratically with the mean.

### Differential Expression Analysis

Differential expression analyses were performed using DESeq2 (v1.38.3) ^36^. We used the ‘pseudobulk’ approach whereby, for a given individual, the read counts for each gene were pooled across all cells in the cluster of interest. By aggregating counts to the cluster level in this way, the data can be treated effectively as bulk and analysed using conventional DE workflows for bulk data. The effect of exposure was assessed while accounting for batch as a covariate. In addition, we evaluated the effects of origin and the origin-by-exposure interaction to identify origin-dependent differences in the transcriptional response to exposure. To determine whether exposure altered baseline origin-associated variation, origin effects were also examined separately within exposed and non-exposed individuals.

As first negative control, individuals were partitioned into two artificial groups (X and Y) containing approximately equal proportions of exposed and non-exposed samples. A second negative control was generated by randomly assigning individuals to two groups (Z and W). Differential expression analyses performed on these artificial groupings were used to confirm that the transcriptional differences observed between exposed and non-exposed fish exceeded those expected by chance alone. Examples from the three biggest clusters (B0, N1, N6) are provided in Supplementary Figure 2.

### Functional analysis of DE genes upon infection

A total of 77 genes with at least 2-fold significant (p ≤ 0.05) expression change in a single cluster and/or any significant (p ≤ 0.05) expression change in at least 2 clusters entered functional analyses. They were functionally annotated by retrieving available information for the respective ENSEMBL ID from ensembl, and by searching the Gene Card database by gene name. If no gene name was available and a gene was considered “novel”, stepwise identification included a search for orthologs through ensembl and, if unsuccessful, a blastp search of the longest ensembl protein sequence to retrieve orthologs, and using the ortholog gene name to retrieve Gene Card information. These self-identified gene names are marked with an asterisk throughout. Gene expression networks and interactions were identified using the STRINGdb on ensembl IDs in R, and visualized in Cytoscape. The retrieved gene networks were then manually expanded based on gene descriptions and gene involvement in particular processes. Heatmap visualisation was performed in R using RStudio 2025.09.2+418 and R version 4.5.1 (2025-06-13 ucrt) Great Square Root on asinh transformed log2 fold change data, using dist and hclust, masking genes and clusters with less than 20 reads in both non-exposed and exposed conditions from the heatmap.

Unsupervised clustering of fold changes for the 77 differentially expressed genes was performed using a combination of distance methods (euclidean, maximum, manhattan, canberra, minkowski) and clustering methods (ward.D, ward.D2, single, complete, average, mcquitty) in R using RStudio 2025.09.2+418 and R version 4.5.0 (2025-04-11 ucrt) How About a Twenty-Six on asinh transformed data. Network analyses were performed using the biomaRt ^37, 38^ and STRINGdb ^39^ packages in R using RStudio 2025.09.2+418 and R version 4.5.0 (2025-04-11 ucrt) How About a Twenty-Six and visualized in Cytoscape 3.10.4 ^40^ and Adobe Illustrator.

### Functional analysis of DE genes by origin

A total of 1104 (in control animals) / 42 (in exposed animals) genes with at least 2-fold significant (p ≤ 0.05) expression change in a single cluster and/or any significant (p ≤ 0.05) expression change in at least 2 clusters entered functional analysis using ShinyGO.

## Results

To understand the genetic relatedness among and within the groups, we called SNPs on transcriptome data and performed kinship analyses. We found that the three origin groups and the treatment groups were genetically indistinguishable. Heatmap, clustering, and principal component analyses of SNPs called from the scRNA-seq data revealed no genetic population structure among the 18 individuals included in the study (Supplementary Figure 3). Any observed gene expression differences were therefore unlikely to result from inadvertent sampling of siblings or from genome-wide allelic divergence among origins.

To assign a fate to each sequenced cell, we performed Seurat clustering. This recovered 30 cell types. Based on conserved vertebrate markers and previously established trout cell-type markers^16^ (Supplementary Table 3; Supplementary Figure 4A), 24 of the 30 Seurat clusters were assigned to specific hematopoietic lineages. These included natural killer cells; seven B-cell clusters, two of which represented pro-/pre-B cells; seven neutrophil clusters; macrophages/monocytes; thrombocytes; two red blood cell clusters; and progenitor cells (Figure 2). B cells and neutrophils were the most abundant cell types, accounting for 36% and 43% of all cells, respectively (Figure 2; Supplementary Table 4). Cluster BN19 was classified as a putative phagocytic B-cell population based on the combined expression of B-cell and neutrophil markers.

**Figure 2.**
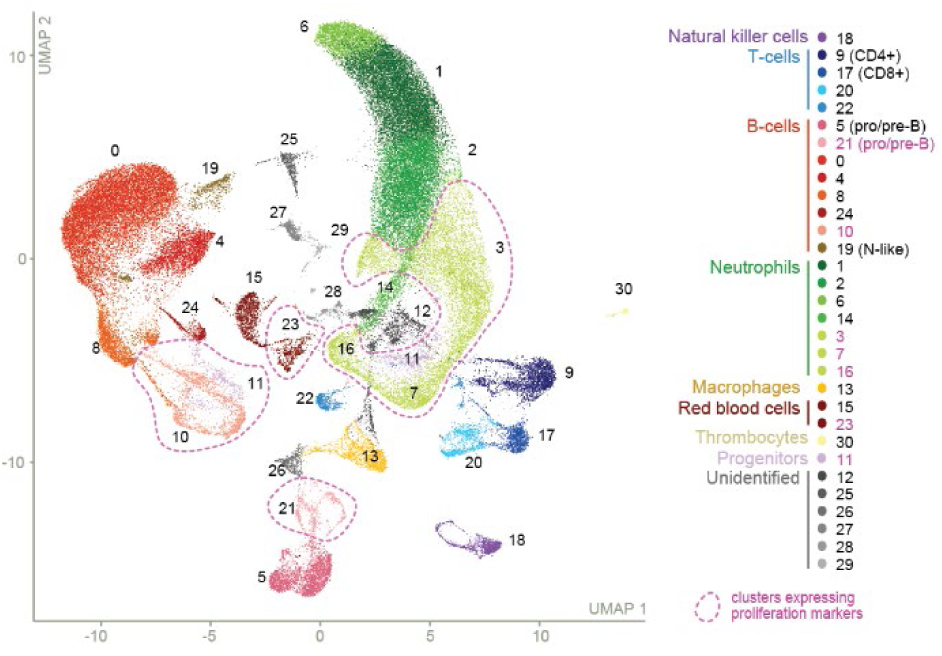
UMAP projection of all cells from all samples. Of 30 clusters identified by Seurat, 24 could be assigned to hematopoietic lineages or subtypes (see legend). Cluster numbers correspond to Seurat-derived cluster IDs. Clusters enriched for proliferation-associated gene expression are highlighted / circled in purple.

Six clusters could not be assigned to a specific immune cell type. NA12 and NA25–NA29 lacked major lineage markers, and their identities could therefore only be inferred tentatively from the expression of selected genes. NA26 may contain hematopoietic stem or progenitor cells, based on the expression of *msi2*, *kita*, *myb*, and *cdk6*, and may include megakaryocyte progenitors expressing *mpl*. NA27 likely represents epithelial or basal epithelial-like cells, indicated by *krt18a.1* and *krt5*, together with high expression of metabolic genes such as *cox6b1, cox7a1, atp5mc3, atp5if1, mdh1*, and *mdh2*. NA28 expressed *scgn* and *atp2a3*, together with *kcnab2a*, *s100a10a*, and *ahnak*, suggesting that it contains at least some neuroendocrine-like cells. Finally, the small NA29 cluster may represent doublets containing stromal cells, marked by *cxcl12*, and macrophages, marked by *mrc1* and *cfd*, potentially with phagocytosed epithelial material expressing *krt18a.1* and *krt5*. Also, exposure-induced gene expression changes were limited in the unassigned clusters. 15 genes with >2 fold up/downregulation upon exposure in other clusters were only lowly expressed and/or hardly responded to exposure in the unassigned clusters, except for NA26 (Supplementary Figure 5). The unassigned clusters were therefore not investigated further.

To understand which clusters contained proliferating cell populations, we specifically analysed expression of four proliferation markers *h2az*, *pcna*, *pclaf*, and *mki67*. Proliferation signaturs were found in clusters B21 (pro- /pre-B cells), B10, N3, N7, N16, RBC23, and P11 (Figure 2; Supplementary Figure 6). Some of these clusters may represent proliferating progenitor populations. For example, based on its co-regulation with both B- and T-cell gene-expression programs described below, P11 may represent a proliferating lymphoid precursor population. Additional clusters, including NK18, expressed proliferation markers only in subsets of cells (Supplementary Figure 6).

To expand the currently limited set of cell-specific marker genes in brown trout, we verified the specificity and prevalence of previously proposed markers. Essentially, markers proposed by Ord et al. (2026)^16^ were screened for lineage-specific and prevalent expression (at least 60% of the cells in a cluster) in this study. The identified genes covered major immune populations and subtypes, including T cells, B cells, neutrophils, and macrophages (examples are presented in Figure 3). A complete overview of the marker genes, their lineage specificity, and their prevalence is provided in Supplementary Table 5, their expression patterns are shown in Supplementary Figure 4B.

**Figure 3.**
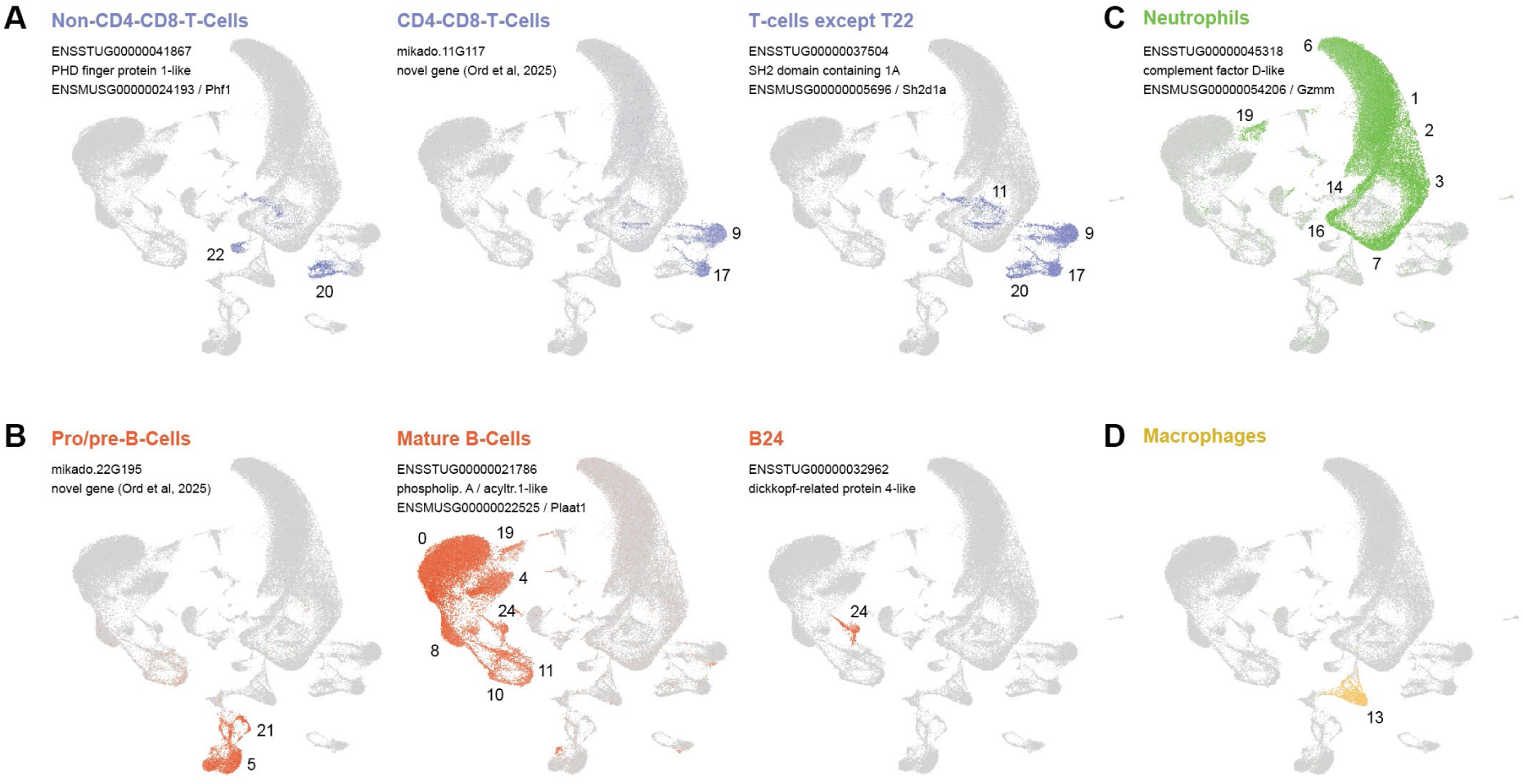
UMAPs highlighting candidate marker genes for major immune-cell populations. UMAP projections show the expression of candidate marker genes for (A) T-cell subsets; (B) pro/pre-B cells, mature B cells, and cluster B24 which may represent naïve B-cells; (C) neutrophils; and (D) macrophages. Cells expressing the marker genes are colored according to the respective immune-cell lineage, all remaining cells are shown in grey. Numbers indicate cell-cluster identities. Above each UMAP, gene identifiers, annotations, and mouse orthologues are provided (if available).

To understand how the trout immune system reacts to PKD exposure, we compared cell abundances and gene expression between exposed and control animals. In terms of cell numbers, a model accounting for origin showed that T-cell abundance increased, while B cells showed mild decreases, neutrophils responded in a cluster-specific manner, and macrophages, RBCs and thrombocytes increased in response to exposure (Figure 4A, Supplementary Figure 7A). In terms of gene expression, differential regulation was particularly prominent in B-cells, neutrophils, and progenitor cells, while non-immune cells (thrombocytes, red blood cells, NA clusters) did not alter gene expression in response to exposure (Figure 4B). 77 genes were significantly (p.adj <= 0.05) regulated >= 2 fold (log2fc) in one cluster, or at any fold change in several clusters (Supplementary Table 6). In line with the sub-clinical infection scenario, gene expression changes were mild (below 10-fold; Figure 4B, Supplementary Figure 7B), so we tested for overarching patterns. We found that fold-change values were sufficient to sort cell types into four major lineages (Neutrophil clusters, T-cells plus NK cells, B-cells, Non-immune cells) by unsupervised clustering (Supplementary Figure 7C), suggesting biological relevance.

**Figure 4.**
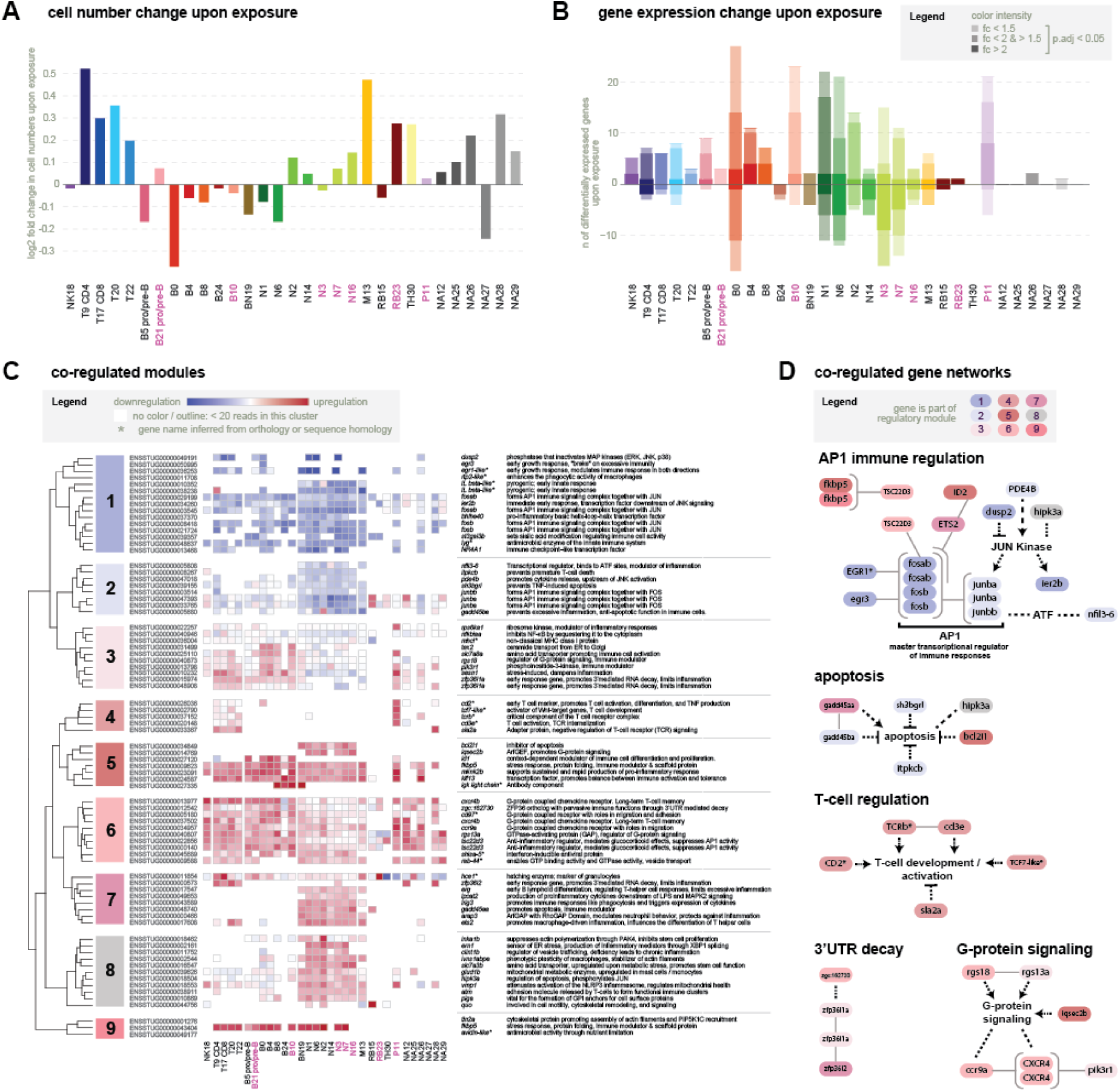
PKD exposure alters relative cell-type abundance and cell-type-specific gene expression. **(A) Bar plot** of log₂ fold change in relative cell abundance for each cell cluster in exposed versus non-exposed fish. Fold changes were estimated using a generalized linear model accounting for origin and experimental batch. Positive values indicate higher relative abundance after exposure. Pink labels indicate clusters with a proliferation signature. **(B) Bar plot** of the number of genes significantly differentially expressed between exposed and non-exposed fish within each cluster (adjusted P < 0.05). Bars above and below zero indicate upregulated and downregulated genes, respectively. Color intensity indicates absolute fold change: dark, >2-fold; intermediate, 1.5–2-fold; and light, <1.5-fold. Pink labels indicate clusters with a proliferation signature. **(C) Heatmap** of expression fold changes for 77 exposure-responsive genes that are either regulated 2-fold in one cluster, or at any level in multiple clusters. The heatmap is based on unsupervised hierarchical clustering using Canberra distance and Ward.D2 linkage. Nine co-regulated modules emerge across cell clusters. Cells with fewer than 20 reads for the respective gene and cluster are left blank. Asterisks indicate gene names inferred from orthology or sequence homology. **(D) Functional association networks** represented among the 77 exposure-responsive genes. Node colors indicate module membership. Grey edges denote associations retrieved through STRINGdb, black dashed edges indicate manually curated relationships based on established gene functions. The displayed networks are associated with AP-1-centred immune regulation, apoptosis, T-cell development and activation, 3′-UTR-mediated mRNA decay, and G-protein-coupled signaling.

To understand which processes reacted to PKD exposure, we performed unsupervised clustering of genes into modules based on fold change regulation. We identified nine cell-type-specific regulatory modules (Figure 4C). These modules represented coordinated immunoregulatory programs (Figure 4D; Supplementary Table 6).

Modules 1 and 2 were broadly expressed across cell types, and predominantly downregulated. They contained immediate-early transcripts associated with AP-1 and EGR signaling, including *fosb/fosab*, *junba/junbb*, *egr1/egr3*, and *ier2b*, together with *bhlhe40* and an *il1b*-like transcript. Negative-feedback regulators such as *dusp2*, *nfil3*, *gadd45ba*, *itpkcb*, and *nr4a1* were also represented. Their coordinated downregulation indicates reduced activity of early MAPK-, calcium-, and stress-responsive transcriptional programs.

Module 3 was upregulated in lymphocytes but downregulated in neutrophils. It included transcripts associated with attenuation of inflammatory signaling, such as *rgs18*, *nfkbiaa*, and *zfp36l1a*, as well as components of PI3K/MAPK signaling and metabolic stress responses. This module therefore showed opposing regulation between lymphoid and myeloid populations.

Module 4 was expressed in T cells and the proliferating P11 cluster. Upregulation of T-cell-associated transcripts, including *cd3e*, a *tcrb*-like gene, *cd2*-like, and *tcf7*-like, in P11 supports its interpretation as a proliferating T-cell precursor population. In contrast, the negative TCR regulator *sla2a* was upregulated in mature T cells but not in P11.

Modules 5 and 6 contained transcripts upregulated broadly across immune cells but not expressed beyond immune cells. These included B-cell-associated transcripts such as *id1* and an immunoglobulin light-chain gene, neutrophil-associated *bcl2l1* and *iqsec2b*, and broadly induced stress and regulatory genes including *mknk2b*, *fkbp5*, *klf13*, *tsc22d3*, and a *zfp36*-like gene. Chemokine receptors and trafficking genes, including *cxcr4b*, *ccr9a*, *cd97*-like, and *rab44*-like, further indicated coordinated changes in cellular positioning and vesicle dynamics. Together, these modules reflected broad immune activation accompanied by regulatory and homeostatic transcriptional programs.

Modules 7 and 8 were predominantly expressed and upregulated in neutrophils. They included transcriptional regulators, cytoskeletal and trafficking genes, and transcripts associated with autophagy, cellular stress, and metabolism. Representative genes included *ets2*, *erg*, *ern1*, *vmp1*, *clint1b*, *lpcat2*, *quo*, *inka1b*, *arap3*, *slc7a3b*, and *glud1b*. Negative regulators such as *zfp36l2* and *gadd45aa* were also present. These patterns were consistent with activated, migratory, and metabolically adapted neutrophils whose inflammatory activity remained transcriptionally regulated.

Module 9 contained mostly low-abundance transcripts, with the notable exception of *fkbp5*, which was strongly upregulated across most immune-cell clusters. A second *fkbp5*-annotated gene in Module 5 showed a similarly broad response.

Across modules, the downregulation of AP-1-associated immediate-early transcripts and the induction of glucocorticoid-responsive and immunoregulatory genes suggest a system-wide transition from early activation towards a more controlled immune state by day 25.

To understand the impact of origin, we first analysed origin-dependence in gene expression across all samples. We found that 2175 genes were significantly differentially expressed between origins when not accounting for treatment, predominantly between wild and farm fish, with the largest effects observed in B-cell populations (Figure 5A). 56 genes were regulated >= 2 fold in response to origin (Supplementary Table 7). These origin-dependent changes were distinct from exposure-dependent changes. While 18 genes were shared between the origin-associated set (n = 56) and the exposure-responsive set (n = 77), these sets were regulated in different cell types: origin-dependent regulation occurred primarily in B cells, whereas exposure-responsive genes were most prevalent in neutrophils and progenitor populations (Figure 5B).

**Figure 5:**
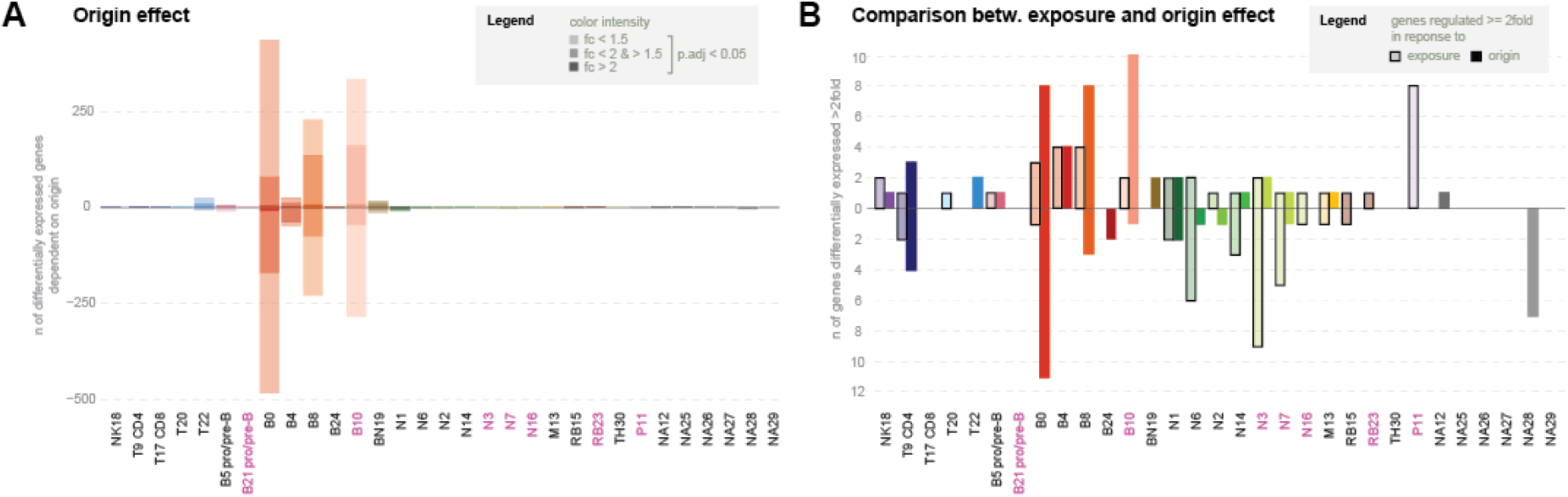
Origin-dependent gene regulation is concentrated in B cells and differs from exposure-dependent gene regulation. **(A) Bar plot** of number of genes with significant origin-dependent expression in each cell cluster. Pairwise comparisons were made between wild and farmed fish and between wild and wild:farm fish. Only genes with an adjusted P value < 0.05 are shown. Color intensity indicates effect size: dark, >2-fold; intermediate, 1.5–2-fold; and light, <1.5-fold. Pink labels indicate proliferating clusters. **(B) Bar plot** of number of genes regulated significantly at least twofold upon exposure (filled bars) or in dependence of origin (outlined bars). Origin-dependent regulation occurred predominantly in B cells, while exposure-responsive genes were concentrated in neutrophil and progenitor populations.

To understand the impact of origin during an active pathogen exposure, we then re-evaluated origin-dependence of gene expression in non-exposed and exposed fish separately. This analysis confirmed the previously observed strong effect of origin, and additionally revealed that this effect was restricted to control animals: 3192 genes were significantly regulated (p.adj <= 0.05; >= 2 fold in one cluster, or at any fold change in several clusters) in controls. In exposed animals, only 89 genes were regulated in a similar fashion (Figure 6A). In the B0 cluster, which is a cluster featuring both exposure-dependent and origin-dependent gene regulation (Figure 4B, Figure 5A), non-exposed samples occupied a broad transcriptional space and separated according to origin, whereas exposed samples from all three origins occupied a more restricted and overlapping region of the PCA space (Figure 6B). Thus, PKD exposure induced a largely shared expression state across origins and eliminated origin-dependent baseline differences in gene expression.

**Figure 6:**
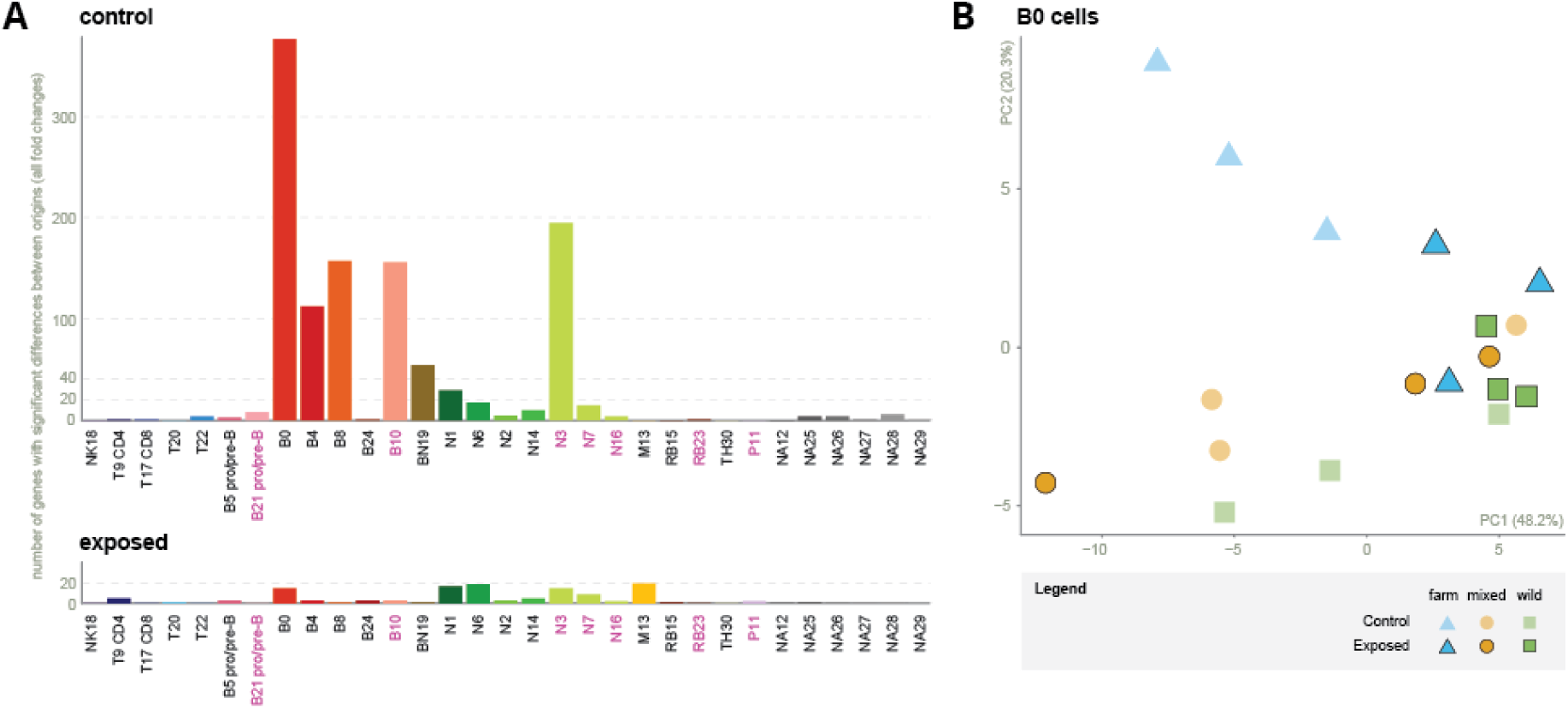
Origin-dependent transcriptional differences diminish following PKD exposure. **(A) Bar plot** of number of genes showing significant origin-dependent expression in each cell cluster under non-exposed conditions (top) and following T. bryosalmonae exposure (bottom).Genes were counted if they were significant (p.adj <= 0.05), and either regulated more than log2fold of 2 / -2, or regulated significantly in multiple clusters. Origin effects were pronounced in non-exposed fish, particularly in B-cell clusters, but were strongly reduced after exposure (n=119). **(B) Principal component analysis** of gene expression in the B0 cluster. Non-exposed samples occupy a broad transcriptional space and show separation among origins. In contrast, exposed samples converge along PC1 and PC2, indicating a shared exposure-associated transcriptional state. Non-exposed samples are shown in light shades, exposed samples in darker colors with black outlines.

## Discussion

In this study, we measured the impact of early life origin and parental living conditions on gene expression in immune cells with and without a pathogen challenge using single cell RNA sequencing. We find a significant transcriptional imprint of origin and early-life conditions on gene expression in immune cells at baseline. Several immune-cell populations, most prominently B cells, showed origin-dependent transcriptional differences. This imprint affects thousands of genes yet did not translate into an origin-dependent response to PKD exposure.

Instead, the imprint disappeared in animals exposed to the pathogen. This finding is notable because wild and hatchery-associated fish have previously been suggested to differ in immune state or immune priming ^3, 41–43, 20^. The result also differs from many studies in humans, where immune stimulation often reveals regulatory differences that are not apparent at baseline ^44–48^. Population-specific responses to infection have likewise been reported in stickleback ^49^. In these systems, immune challenge amplifies underlying genetic or regulatory variation rather than reducing it.

Our data, however, contradicts this expectation. We found no evidence that origin modified the immune response to *T. bryosalmonae* exposure. In contrast, we observe a canalization of distinct immune cell gene expression during the pathogen challenge, a loss of origin signatures, and a common response. This parallels finding im wild field voles, where a subset of genes with high interindividual variation at baseline became less variable after standardized immune stimulation, although other genes showed the opposite pattern ^50^. Shared transcriptional responses have also been observed in stickleback transplanted between lakes, where immune expression shifted towards the profile of the destination environment ^51^, across divergent *Drosophila* species and lines following parasitoid attack ^52^, and in genetically resistant and susceptible rainbow trout following viral infection ^53^. Together with this evidence, our study suggests that strong environmental or immune challenges can drive divergent backgrounds towards a common functional state.

In terms of the presence of cell types and their abundance in control and exposed animals, our data is consistent with previous studies. Under control conditions, B cells and neutrophils dominated the immune-cell composition of the brown trout kidney ^54, 31, 16, 17^. One cluster co-expressing canonical B-cell and neutrophil markers likely corresponds to the phagocytic B-cell population previously designated NA2 ^16^. These cells combine antibody-mediated functions with phagocytic capacity and may therefore provide an important functional link between innate and adaptive immunity ^55^. Phagocytic B cells illustrate how comparative studies in non-model vertebrates can reveal previously unknown immune functions, as they were first described in fish, amphibians, and reptiles ^56–59^ and only subsequently confirmed in mammals ^60, 61^. Under exposed conditions, we find a relative increase in T cells and in putative non-immune populations. These shifts represent changes in relative cell frequencies rather than absolute expansion or depletion of individual populations, and are consistent with tissue remodelling in the kidney. They are also broadly consistent with previous descriptions of renal lymphoid hyperplasia, altered myeloid activity, dysregulated B-cell and immunoglobulin responses, and T-helper-like transcriptional signatures in PKD ^62, 63, 25, 27, 28^ and with previous antibody-based analyses in the same experimental system ^17^.

The most prominent transcriptional response to pathogen exposure occurred in B cells, neutrophils, and proliferating progenitor populations. Thus, the cell types showing the largest compositional shifts were not necessarily those undergoing the strongest transcriptional regulation. This distinction highlights the value of jointly assessing cell frequencies and cell-type-specific gene expression: a population may respond through changes in abundance, transcriptional state, or both. We found no clear evidence for a previously reported broadly polarized Th2 response ^25^. However, increased *klf13* expression in T cells may indicate enhanced Th2-associated effector functions, including regulation of IL-4 expression (Kwon et al. 2014). This interpretation remains tentative because it is based on a limited number of markers.

Within the transcriptional response to PKD exposure, a prominent feature was the downregulation of immediate-early and AP-1-associated genes, including members of the *jun* and *fos* families. Together with the opposing responses of innate and adaptive compartments, this pattern suggests a transition away from broad early activation towards a more regulated immune state. The transcriptomic profile appears to reflect an active but restrained response to infection, which is in line with the previously reported course of infection in this experimental population, devoid of macroscopic PKD lesions or exposure-associated mortality ^17^.

Histopathology was not performed on the specific individuals used for scRNA-seq because of limited tissue availability, and mild infection-associated lesions therefore cannot be excluded. Nevertheless, the combined clinical and transcriptomic evidence indicates that the dataset primarily captures a subclinical stage of infection.

PKD outcomes depend strongly on environmental and host context, including temperature, infection pressure, and water quality ^21, 23^. ^25^ reported a temperature-associated shift from a predominantly Th1-like response at 12°C towards a Th2-like response at 15°C. Our fish were maintained above that threshold (16°C) yet showed only limited evidence for Th2 polarization and no overt disease. Temperature alone is therefore unlikely to determine whether infection remains controlled or progresses to immunopathology. Instead, disease severity probably emerges from interactions among temperature, parasite burden, host condition, and the regulation of specific immune pathways.

In the future, the expression programs identified in this study may help distinguish controlled infection from progression towards overexpressed inflammation and renal pathology. Their comparison across environmental conditions, trout populations, and fish species may clarify why salmonids develop severe PKD in some cases, whereas under different conditions salmonids and other freshwater fishes sustain active infections and parasite excretion without overt disease ^17, 64^. Such analyses could identify the points at which immune control fails and uncover factors that determine whether infection results in tolerance or clinical disease.

These future studies could largely benefit from the novel markers confirmed in this study. Several pan-lineage and subset-specific markers for B cells, T cells, neutrophils, and macrophages ^16^ could be confirmed, which substantially expands the available molecular toolkit for immune-cell identification in salmonids. At present, flow-cytometric reagents in brown trout are largely restricted to IgM+ B cells, myeloid cells, and CD8+ T cells ^17^, with similarly limited coverage in rainbow trout ^65^. Given the ecological and economic importance of salmonids, a broader marker set will facilitate more detailed monitoring of immune-cell composition and activation. Also, our expression data suggest that existing antibodies only partially label the desired cell populations (**Supplementary Figure 6**). The candidate markers now require validation by qPCR in sorted or enriched cell populations. Once validated, they could be applied across life stages, tissues, and pathogen-exposure conditions. Markers encoding cell-surface proteins may additionally support the development and validation of new antibodies, further extending the experimental toolkit for salmonid immunology.

## Conclusion

Our detailed single-cell data of the brown trout immune system in animals exposed to PKD show no evidence for an origin-dependent reaction pattern towards a subclinical infection with *T. bryosalmonae*. Thus, other explanations for a poor performance of farmed fish in the wild, such as a stimulus deprived environment ^66, 67^ or inappropriate predator responses ^68^, may need to be explored. Also, hatchery–wild differences regarding immune responses may emerge under more severe or ecologically complex exposure conditions. This would require similar analyses in longitudinal, field-based samplings or lab experiments using higher infection doses, co-infections, or other challenges.

## Author contributions

Conceptualization: HSP, IAK Design: HSM, HSP, IAK

Methodology: HSM, JO, QC, ST, ND, HSP, IAK

Data collection: HSM, JO

Data curation: JO, QC

Validation: QC

Formal analysis: QC, ST, HSP, IAK

Resources: HSP, IAK

Writing – figure preparation: QC, IAK

Writing – original draft preparation: QC, ST, HSP, IAK

Writing – review and editing: QC, HSM, JO, ND, ST, HSP, IAK

## Acknowledgements

We are grateful to Helmut Segner for critical reading of the draft manuscript, Gary Delalay for feedback on statistics, Simone Oberhänsli for discussion of data analysis strategies, and the EpiEvo group at FIWI for discussion of data, figures and results. We thank the Fish Wardens of the Canton of St Gallen and the staff and management of Fischereizentrum Steinach (Canton of St Gallen) for assistance with fish acquisition, Pamela Nicholson of the University of Bern, Next Generation Sequencing Platform for advice regarding sample preparation and for performing 10x Chromium sequencing, and Robine Schoch and Sarah Posthaus for support with trout husbandry.

## Permits

Experiments were performed under permit # 33570 (BE25/2021) from the Veterinary Office of the Canton of Bern, Switzerland.

## Data availability

Raw data used for analysis are available under accession PRJEB113617 in the European Nucleotide Archive (ENA). Scripts used to analyse the data are available at https://github.com/FIWI-UniBe/TRIP-scRNA-Analysis-Scripts.

## Declaration of generative AI and AI-assisted technologies in the writing process

During the preparation of this work, the authors used ChatGPT Plus in order to improve the language and readability of the article. After using this tool/service, the authors reviewed and edited the content as needed and take full responsibility for the content of the published article.

## References

1. Gollwitzer, E.S., and Marsland, B.J. (2015). Impact of Early-Life Exposures on Immune Maturation and Susceptibility to Disease. Trends in immunology 36, 684–696. 10.1016/j.it.2015.09.009.

2. Roth, O., Beemelmanns, A., Barribeau, S.M., and Sadd, B.M. (2018). Recent advances in vertebrate and invertebrate transgenerational immunity in the light of ecology and evolution. Heredity 121, 225–238. 10.1038/s41437-018-0101-2.

3. Karvonen, A., Aalto-Araneda, M., Virtala, A.-M., Kortet, R., Koski, P., and Hyvärinen, P. (2016). Enriched rearing environment and wild genetic background can enhance survival and disease resistance of salmonid fishes during parasite epidemics. Journal of Applied Ecology 53, 213–221. 10.1111/1365-2664.12568.

4. Bonduriansky, R., and Day, T. (2009). Nongenetic Inheritance and Its Evolutionary Implications. Annu. Rev. Ecol. Evol. Syst. 40, 103–125. 10.1146/annurev.ecolsys.39.110707.173441.

5. Badyaev, A.V., and Uller, T. (2009). Parental effects in ecology and evolution: mechanisms, processes and implications. Philosophical transactions of the Royal Society of London. Series B, Biological sciences 364, 1169–1177. 10.1098/rstb.2008.0302.

6. Andreou, D., Arkush, K.D., Guégan, J.-F., and Gozlan, R.E. (2012). Introduced pathogens and native freshwater biodiversity: a case study of Sphaerothecum destruens. PloS one 7, e36998. 10.1371/journal.pone.0036998.

7. Gozlan, R.E., St-Hilaire, S., Feist, S.W., Martin, P., and Kent, M.L. (2005). Biodiversity: disease threat to European fish. Nature 435, 1046. 10.1038/4351046a.

8. FAO (2024). The State of World Fisheries and Aquaculture 2024 (FAO).

9. BAFU (2026). Federal stocking statistics. https://www.fischereistatistik.ch/.

10. Aurélie, R., and Rubin, J.-F. (2026). Response of a Brown Trout Population to Anthropic and Environmental Factors Over 25 Years in the Boiron of Morges, Switzerland. Fisheries Management Eco 33, 40–53. 10.1111/fme.70006.

11. Caudron, A., Champigneulle, A., Largiadèr, C.R., Launey, S., and Guyomard, R. (2009). Stocking of native Mediterranean brown trout (Salmo trutta) into French tributaries of Lake Geneva does not contribute to lake-migratory spawners. Ecology of Freshwater Fish 18, 585–593. 10.1111/j.1600-0633.2009.00374.x.

12. Champigneulle, A., Largiadèr, C.R., and Caudron, A. (2003). REPRODUCTION DE LA TRUITE (Salmo trutta L.) DANS LETORRENT DE CHEVENNE, HAUTE-SAVOIE.UN FONCTIONNEMENT ORIGINAL ? Bull. Fr. Pêche Piscic., 41–70. 10.1051/kmae:2003021.

13. Bowden, T.J. (2008). Modulation of the immune system of fish by their environment. Fish & shellfish immunology 25, 373–383. 10.1016/j.fsi.2008.03.017.

14. Christie, M.R., Marine, M.L., Fox, S.E., French, R.A., and Blouin, M.S. (2016). A single generation of domestication heritably alters the expression of hundreds of genes. Nature communications 7, 10676. 10.1038/ncomms10676.

15. Rehberger, K., Wernicke von Siebenthal, E., Bailey, C., Bregy, P., Fasel, M., Herzog, E.L., Neumann, S., Schmidt-Posthaus, H., and Segner, H. (2020). Long-term exposure to low 17α-ethinylestradiol (EE2) concentrations disrupts both the reproductive and the immune system of juvenile rainbow trout, Oncorhynchus mykiss. Environment international 142, 105836. 10.1016/j.envint.2020.105836.

16. Ord, J., Martinez, H.S., Solbakken, M.H., Berezenko, A., Oberhaensli, S., Talker, S., Schmidt-Posthaus, H., and Adrian-Kalchhauser, I. (2026). Single-cell analysis of a salmonid immune system (river brown trout Salmo trutta fario) reveals evolutionary divergence and hatchery-induced transcriptional reprogramming. BMC biology 24. 10.1186/s12915-026-02554-2.

17. Saura Martinez, H., Delalay, G., Talker, S., and Schmidt-Posthaus, H. (2025). Parental influence on brown trout offspring immune cell composition: An infection study with Tetracapsuloides bryosalmonae. PloS one 20, e0308779. 10.1371/journal.pone.0308779.

18. Anderson, C.L., Canning, E.U., and Okamura, B. (1999). Molecular data implicate bryozoans as hosts for PKX (phylum Myxozoa) and identify a clade of bryozoan parasites within the Myxozoa. Parasitology 119 (Pt 6), 555–561. 10.1017/s003118209900520x.

19. Okamura, B., Anderson, C.L., Longshaw, M., Feist, S.W., and Canning, E.U. (2001). PATTERNS OF OCCURRENCE AND 18S rDNA SEQUENCE VARIATION OF PKX (TETRACAPSULA BRYOSALMONAE), THE CAUSATIVE AGENT OF SALMONID PROLIFERATIVE KIDNEY DISEASE. Journal of Parasitology 87, 379–385. 10.1645/0022-3395(2001)087[0379:POOARS]2.0.CO;2.

20. Strepparava, N., Ros, A., Hartikainen, H., Schmidt-Posthaus, H., Wahli, T., Segner, H., and Bailey, C. (2020). Effects of parasite concentrations on infection dynamics and proliferative kidney disease pathogenesis in brown trout (Salmo trutta). Transboundary and emerging diseases 67, 2642–2652. 10.1111/tbed.13615.

21. Strepparava, N., Segner, H., Ros, A., Hartikainen, H., Schmidt-Posthaus, H., and Wahli, T. (2018). Temperature-related parasite infection dynamics: the case of proliferative kidney disease of brown trout. Parasitology 145, 281–291. 10.1017/S0031182017001482.

22. Bettge, K., Wahli, T., Segner, H., and Schmidt-Posthaus, H. (2009). Proliferative kidney disease in rainbow trout: time- and temperature-related renal pathology and parasite distribution. Diseases of aquatic organisms 83, 67–76. 10.3354/dao01989.

23. Saura Martinez, H., Egloff, N., and Schmidt-Posthaus, H. (2023). Investigation of Proliferative Kidney Disease in Brown Trout and Habitat Characteristics Associated with a Swiss Wastewater Treatment Plant. Environments 10, 152. 10.3390/environments10090152.

24. Rubin, A., Coulon, P. de, Bailey, C., Segner, H., Wahli, T., and Rubin, J.-F. (2019). Keeping an Eye on Wild Brown Trout (Salmo trutta) Populations: Correlation Between Temperature, Environmental Parameters, and Proliferative Kidney Disease. Frontiers in veterinary science 6, 281. 10.3389/fvets.2019.00281.

25. Bailey, C., Segner, H., Casanova-Nakayama, A., and Wahli, T. (2017). Who needs the hotspot? The effect of temperature on the fish host immune response to Tetracapsuloides bryosalmonae the causative agent of proliferative kidney disease. Fish & shellfish immunology 63, 424–437. 10.1016/j.fsi.2017.02.039.

26. Bailey, C., Strepparava, N., Wahli, T., and Segner, H. (2019). Exploring the immune response, tolerance and resistance in proliferative kidney disease of salmonids. Developmental and comparative immunology 90, 165–175. 10.1016/j.dci.2018.09.015.

27. Bailey, C., Holland, J.W., Secombes, C.J., and Tafalla, C. (2020). A portrait of the immune response to proliferative kidney disease (PKD) in rainbow trout. Parasite immunology 42, e12730. 10.1111/pim.12730.

28. Bailey, C., Segner, H., Wahli, T., and Tafalla, C. (2020). Back From the Brink: Alterations in B and T Cell Responses Modulate Recovery of Rainbow Trout From Chronic Immunopathological Tetracapsuloides bryosalmonae Infection. Frontiers in immunology 11, 1093. 10.3389/fimmu.2020.01093.

29. Tang, Q., Iyer, S., Lobbardi, R., Moore, J.C., Chen, H., Lareau, C., Hebert, C., Shaw, M.L., Neftel, C., and Suva, M.L., et al. (2017). Dissecting hematopoietic and renal cell heterogeneity in adult zebrafish at single-cell resolution using RNA sequencing. The Journal of experimental medicine 214, 2875–2887. 10.1084/jem.20170976.

30. Hernández, P.P., Strzelecka, P.M., Athanasiadis, E.I., Hall, D., Robalo, A.F., Collins, C.M., Boudinot, P., Levraud, J.-P., and Cvejic, A. (2018). Single-cell transcriptional analysis reveals ILC-like cells in zebrafish. Science immunology 3. 10.1126/sciimmunol.aau5265.

31. Perdiguero, P., Morel, E., and Tafalla, C. (2021). Diversity of Rainbow Trout Blood B Cells Revealed by Single Cell RNA Sequencing. Biology 10. 10.3390/biology10060511.

32. Taylor, R.S., Ruiz Daniels, R., Dobie, R., Naseer, S., Clark, T.C., Henderson, N.C., Boudinot, P., Martin, S.A.M., and Macqueen, D.J. (2022). Single cell transcriptomics of Atlantic salmon (Salmo salar L.) liver reveals cellular heterogeneity and immunological responses to challenge by Aeromonas salmonicida. Frontiers in immunology 13, 984799. 10.3389/fimmu.2022.984799.

33. Zafar, H., Wang, Y., Nakhleh, L., Navin, N., and Chen, K. (2016). Monovar: single-nucleotide variant detection in single cells. Nature methods 13, 505–507. 10.1038/nmeth.3835.

34. R Core Team (2024). R: A Language and Environment for Statistical Computing. (R Foundation for Statistical Computing).

35. Brooks, M., Kristensen, K., Benthem, K., Magnusson, A., Berg, C., Nielsen, A., Skaug, H., Mächler, M., and Bolker, B. (2017). glmmTMB Balances Speed and Flexibility Among Packages for Zero-inflated Generalized Linear Mixed Modeling. The R Journal 9, 378. 10.32614/RJ-2017-066.

36. Love, M.I., Huber, W., and Anders, S. (2014). Moderated estimation of fold change and dispersion for RNA-seq data with DESeq2. Genome biology 15, 550. 10.1186/s13059-014-0550-8.

37. Durinck, S., Moreau, Y., Kasprzyk, A., Davis, S., Moor, B. de, Brazma, A., and Huber, W. (2005). BioMart and Bioconductor: a powerful link between biological databases and microarray data analysis. Bioinformatics (Oxford, England) 21, 3439–3440. 10.1093/bioinformatics/bti525.

38. Durinck, S., Spellman, P.T., Birney, E., and Huber, W. (2009). Mapping identifiers for the integration of genomic datasets with the R/Bioconductor package biomaRt. Nature protocols 4, 1184–1191. 10.1038/nprot.2009.97.

39. Szklarczyk, D., Kirsch, R., Koutrouli, M., Nastou, K., Mehryary, F., Hachilif, R., Gable, A.L., Fang, T., Doncheva, N.T., and Pyysalo, S., et al. (2023). The STRING database in 2023: protein-protein association networks and functional enrichment analyses for any sequenced genome of interest. Nucleic acids research 51, D638–D646. 10.1093/nar/gkac1000.

40. Shannon, P., Markiel, A., Ozier, O., Baliga, N.S., Wang, J.T., Ramage, D., Amin, N., Schwikowski, B., and Ideker, T. (2003). Cytoscape: a software environment for integrated models of biomolecular interaction networks. Genome research 13, 2498–2504. 10.1101/gr.1239303.

41. Nekouei, O., Vanderstichel, R., Kaukinen, K.H., Thakur, K., Ming, T., Patterson, D.A., Trudel, M., Neville, C., and Miller, K.M. (2019). Comparison of infectious agents detected from hatchery and wild juvenile Coho salmon in British Columbia, 2008-2018. PloS one 14, e0221956. 10.1371/journal.pone.0221956.

42. Braden, L.M., Rasmussen, K.J., Purcell, S.L., Ellis, L., Mahony, A., Cho, S., Whyte, S.K., Jones, S.R.M., and Fast, M.D. (2018). Acquired Protective Immunity in Atlantic Salmon Salmo salar against the Myxozoan Kudoa thyrsites Involves Induction of MHIIβ+ CD83+ Antigen-Presenting Cells. Infection and immunity 86. 10.1128/IAI.00556-17.

43. Stølen Ugelvik, M., Mennerat, A., Mæhle, S., and Dalvin, S. (2023). Repeated exposure affects susceptibility and responses of Atlantic salmon (Salmo salar) towards the ectoparasitic salmon lice (Lepeophtheirus salmonis). Parasitology 150, 990–1005. 10.1017/s0031182023000847.

44. Fairfax, B.P., Humburg, P., Makino, S., Naranbhai, V., Wong, D., Lau, E., Jostins, L., Plant, K., Andrews, R., and McGee, C., et al. (2014). Innate immune activity conditions the effect of regulatory variants upon monocyte gene expression. Science (New York, N.Y.) 343, 1246949. 10.1126/science.1246949.

45. Lee, M.N., Ye, C., Villani, A.-C., Raj, T., Li, W., Eisenhaure, T.M., Imboywa, S.H., Chipendo, P.I., Ran, F.A., and Slowikowski, K., et al. (2014). Common genetic variants modulate pathogen-sensing responses in human dendritic cells. Science (New York, N.Y.) 343, 1246980. 10.1126/science.1246980.

46. Aracena, K.A., Lin, Y.-L., Luo, K., Pacis, A., Gona, S., Mu, Z., Yotova, V., Sindeaux, R., Pramatarova, A., and Simon, M.-M., et al. (2024). Epigenetic variation impacts individual differences in the transcriptional response to influenza infection. Nature genetics 56, 408–419. 10.1038/s41588-024-01668-z.

47. Nédélec, Y., Sanz, J., Baharian, G., Szpiech, Z.A., Pacis, A., Dumaine, A., Grenier, J.-C., Freiman, A., Sams, A.J., and Hebert, S., et al. (2016). Genetic Ancestry and Natural Selection Drive Population Differences in Immune Responses to Pathogens. Cell 167, 657–669.e21. 10.1016/j.cell.2016.09.025.

48. Piasecka, B., Duffy, D., Urrutia, A., Quach, H., Patin, E., Posseme, C., Bergstedt, J., Charbit, B., Rouilly, V., and MacPherson, C.R., et al. (2018). Distinctive roles of age, sex, and genetics in shaping transcriptional variation of human immune responses to microbial challenges. Proceedings of the National Academy of Sciences of the United States of America 115, E488–E497. 10.1073/pnas.1714765115.

49. Lohman, B.K., Steinel, N.C., Weber, J.N., and Bolnick, D.I. (2017). Gene Expression Contributes to the Recent Evolution of Host Resistance in a Model Host Parasite System. Frontiers in immunology 8, 1071. 10.3389/fimmu.2017.01071.

50. Wanelik, K.M., Begon, M., Arriero, E., Bradley, J.E., Friberg, I.M., Jackson, J.A., Taylor, C.H., and Paterson, S. (2020). Transcriptome-wide analysis reveals different categories of response to a standardised immune challenge in a wild rodent. Scientific reports 10, 7444. 10.1038/s41598-020-64307-7.

51. Stutz, W.E., Schmerer, M., Coates, J.L., and Bolnick, D.I. (2015). Among-lake reciprocal transplants induce convergent expression of immune genes in threespine stickleback. Molecular ecology 24, 4629–4646. 10.1111/mec.13295.

52. Salazar-Jaramillo, L., Jalvingh, K.M., Haan, A. de, Kraaijeveld, K., Buermans, H., and Wertheim, B. (2017). Inter- and intra-species variation in genome-wide gene expression of Drosophila in response to parasitoid wasp attack. BMC genomics 18, 331. 10.1186/s12864-017-3697-3.

53. Clark, T.C., Thomas, V., Taylor, R.S., Charles, M., Laurent, A., Schwartz-Cornil, I., Collet, B., Lallias, D., Macqueen, D.J., and Martin, S.A.M., et al. (2025). Immune cell-resolved transcriptomics provides insights into the basis for variations of fish genetic resistance to viral disease. BMC biology 23, 348. 10.1186/s12915-025-02452-z.

54. Andresen, A.M.S., Taylor, R.S., Grimholt, U., Daniels, R.R., Sun, J., Dobie, R., Henderson, N.C., Martin, S.A.M., Macqueen, D.J., and Fosse, J.H. (2024). Mapping the cellular landscape of Atlantic salmon head kidney by single cell and single nucleus transcriptomics. Fish & shellfish immunology 146, 109357. 10.1016/j.fsi.2024.109357.

55. Morelli, D.M., Langille, M., Zhang, R., Craig, H.C., Eltom Mohamed, A., Shooshtari, P., Heit, B., and Kerfoot, S.M. (2026). B-cell subsets have different capacities for phagocytosis and subsequent presentation of antigen to cognate T cells. Journal of immunology (Baltimore, Md. : 1950) 215. 10.1093/jimmun/vkaf282.

56. Li, J., Barreda, D.R., Zhang, Y.-A., Boshra, H., Gelman, A.E., Lapatra, S., Tort, L., and Sunyer, J.O. (2006). B lymphocytes from early vertebrates have potent phagocytic and microbicidal abilities. Nature immunology 7, 1116–1124. 10.1038/ni1389.

57. Øverland, H.S., Pettersen, E.F., Rønneseth, A., and Wergeland, H.I. (2010). Phagocytosis by B-cells and neutrophils in Atlantic salmon (Salmo salar L.) and Atlantic cod (Gadus morhua L.). Fish & shellfish immunology 28, 193–204. 10.1016/j.fsi.2009.10.021.

58. Zimmerman, L.M., Vogel, L.A., Edwards, K.A., and Bowden, R.M. (2010). Phagocytic B cells in a reptile. Biology letters 6, 270–273. 10.1098/rsbl.2009.0692.

59. Esteban, M.Á., Cuesta, A., Chaves-Pozo, E., and Meseguer, J. (2015). Phagocytosis in Teleosts. Implications of the New Cells Involved. Biology 4, 907–922. 10.3390/biology4040907.

60. Martínez-Riaño, A., Bovolenta, E.R., Mendoza, P., Oeste, C.L., Martín-Bermejo, M.J., Bovolenta, P., Turner, M., Martínez-Martín, N., and Alarcón, B. (2018). Antigen phagocytosis by B cells is required for a potent humoral response. EMBO reports 19. 10.15252/embr.201846016.

61. Gao, J., Ma, X., Gu, W., Fu, M., An, J., Xing, Y., Gao, T., Li, W., and Liu, Y. (2012). Novel functions of murine B1 cells: active phagocytic and microbicidal abilities. European journal of immunology 42, 982–992. 10.1002/eji.201141519.

62. Abos, B., Estensoro, I., Perdiguero, P., Faber, M., Hu, Y., Díaz Rosales, P., Granja, A.G., Secombes, C.J., Holland, J.W., and Tafalla, C. (2018). Dysregulation of B Cell Activity During Proliferative Kidney Disease in Rainbow Trout. Frontiers in immunology 9, 1203. 10.3389/fimmu.2018.01203.

63. Gorgoglione, B., Wang, T., Secombes, C.J., and Holland, J.W. (2013). Immune gene expression profiling of Proliferative Kidney Disease in rainbow trout Oncorhynchus mykiss reveals a dominance of anti-inflammatory, antibody and T helper cell-like activities. Veterinary research 44, 55. 10.1186/1297-9716-44-55.

64. Möckli, C., Diserens, N., Delalay, G., Moré, G., and Schmidt-Posthaus, H. (2026). Challenging Host Specificity: Malacosporean Parasite Dynamics-Tetracapsuloides bryosalmonae Beyond Salmonids-An Infection Experiment. Journal of fish diseases, e70233. 10.1111/jfd.70233.

65. Korytář, T., Dang Thi, H., Takizawa, F., and Köllner, B. (2013). A multicolour flow cytometry identifying defined leukocyte subsets of rainbow trout (Oncorhynchus mykiss). Fish & shellfish immunology 35, 2017–2019. 10.1016/j.fsi.2013.09.025.

66. Mes, D., van Os, R., Gorissen, M., Ebbesson, L.O.E., Finstad, B., Mayer, I., and Vindas, M.A. (2019). Effects of environmental enrichment on forebrain neural plasticity and survival success of stocked Atlantic salmon. The Journal of experimental biology 222. 10.1242/jeb.212258.

67. Johnsson, J.I., Brockmark, S., and Näslund, J. (2014). Environmental effects on behavioural development consequences for fitness of captive-reared fishes in the wild. Journal of fish biology 85, 1946–1971. 10.1111/jfb.12547.

68. Gro Vea Salvanes, A., and Braithwaite, V. (2006). The need to understand the behaviour of fish reared for mariculture or restocking. ICES Journal of Marine Science 63, 345–354. 10.1016/j.icesjms.2005.11.010.

