## Supplemental Figures for "Early-life conditions shape baseline immune gene expression, but not pathogen-induced immune activation, in brown trout"

**Supplementary Figure 1**

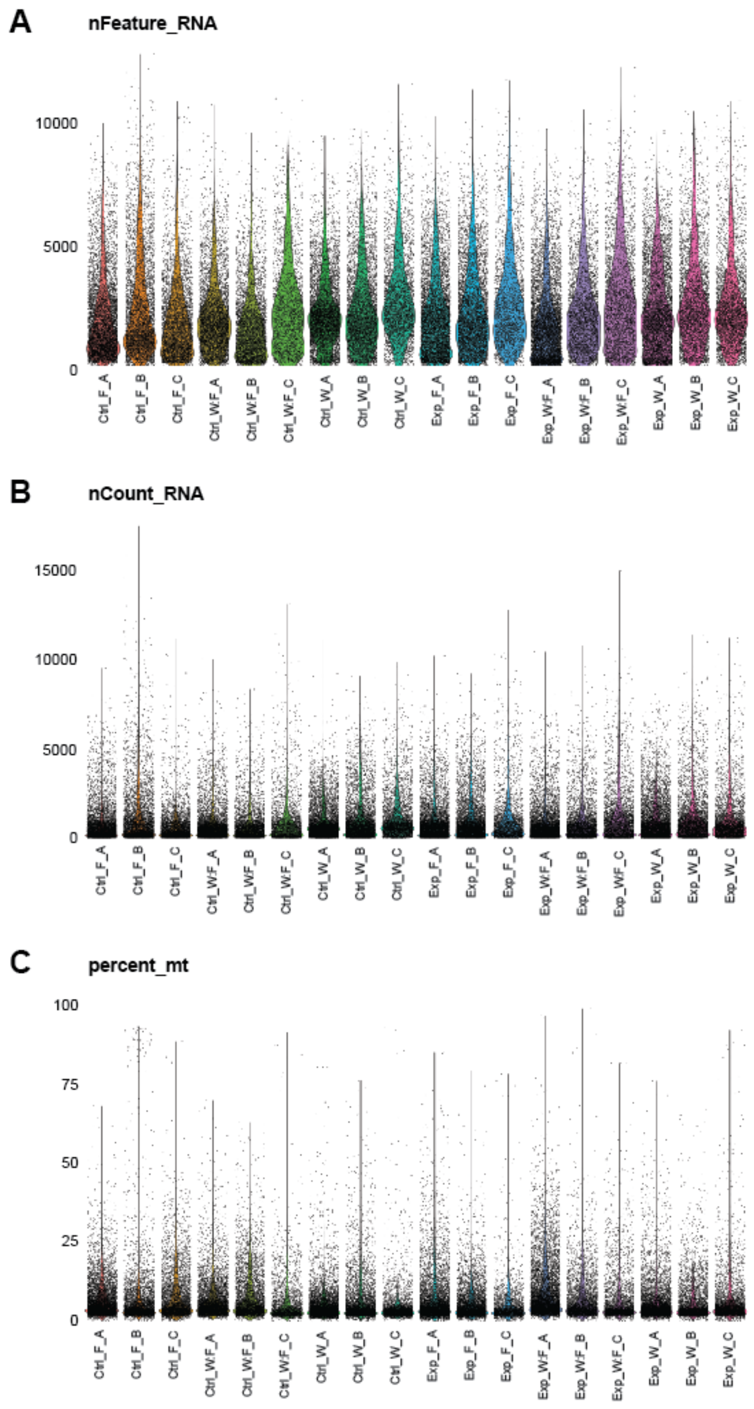


***Supplementary Figure 1. Quality-control metrics for the single-cell RNA-sequencing dataset before filtering.***

*Violin plots show the distribution of (A) the number of genes detected per cell (nFeature_RNA), (B) the total number of RNA counts per cell (nCount_RNA), and (C) the percentage of transcripts assigned to mitochondrial genes (percent_mt) for each sample. Individual points represent single cells, and colours distinguish samples. Sample labels indicate treatment (Ctrl vs Exp), origin (F, W:F, W), and experimental batch (A, B, C).*

### Supplementary Figure 2

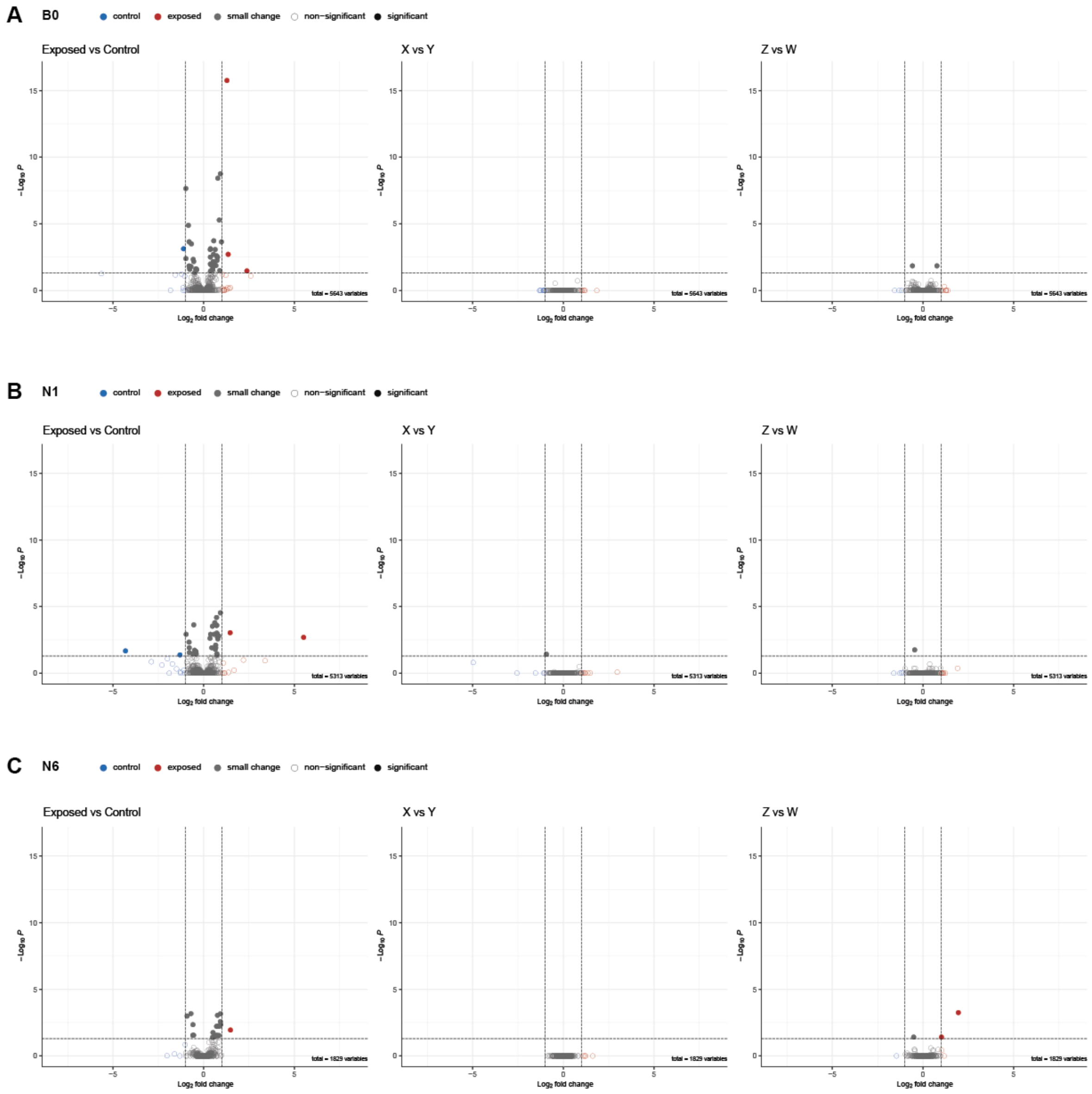


***Supplementary Figure 2. Permutation analysis.***

*Volcano plots show differential expression in (A) B-cell cluster B0, (B) neutrophil cluster N1 and (C) neutrophil cluster N6. For each cluster, columns show the exposed-versus-control comparison in the first panel, and two control contrasts (X versus Y; balanced assortment of control and exposure to each group; and Z versus W; randomized assortment of control and exposure to each group) in the second and third panel respectively. Both permutations (X versus Y and Z versus W) eliminate the differential gene expression observed in the contrast of control and exposure.*

### Supplementary Figure 3

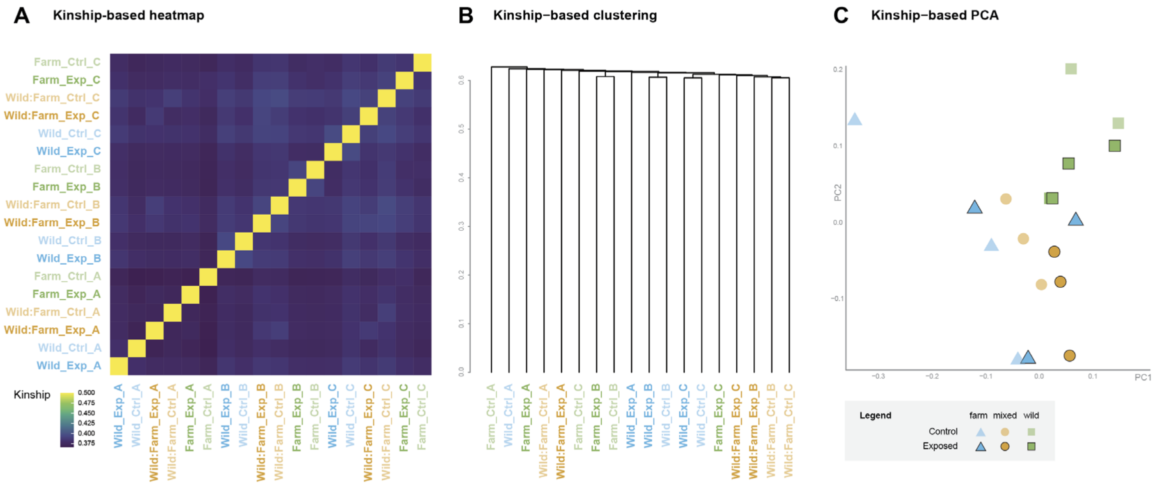


***Supplementary Figure 3. Kinship-based assessment of genetic structure among sequenced individuals.***

*(A) Heatmap of pairwise kinship estimates among the 18 individuals included in the single-cell RNA-sequencing analysis. (B) Hierarchical clustering of individuals based on a distance of 1 - KING Coefficient. (C) Principal component analysis of kinship estimates. Sample labels indicate origin (Wild, Wild:Farm, or Farm), treatment (Ctrl, control; Exp, exposed), and experimental batch (A–C). In the PCA, triangles, circles, and squares represent Wild, Wild:Farm, and Farm individuals, respectively; lighter and darker symbols indicate control and exposed individuals.*

**Supplementary Figure 4**

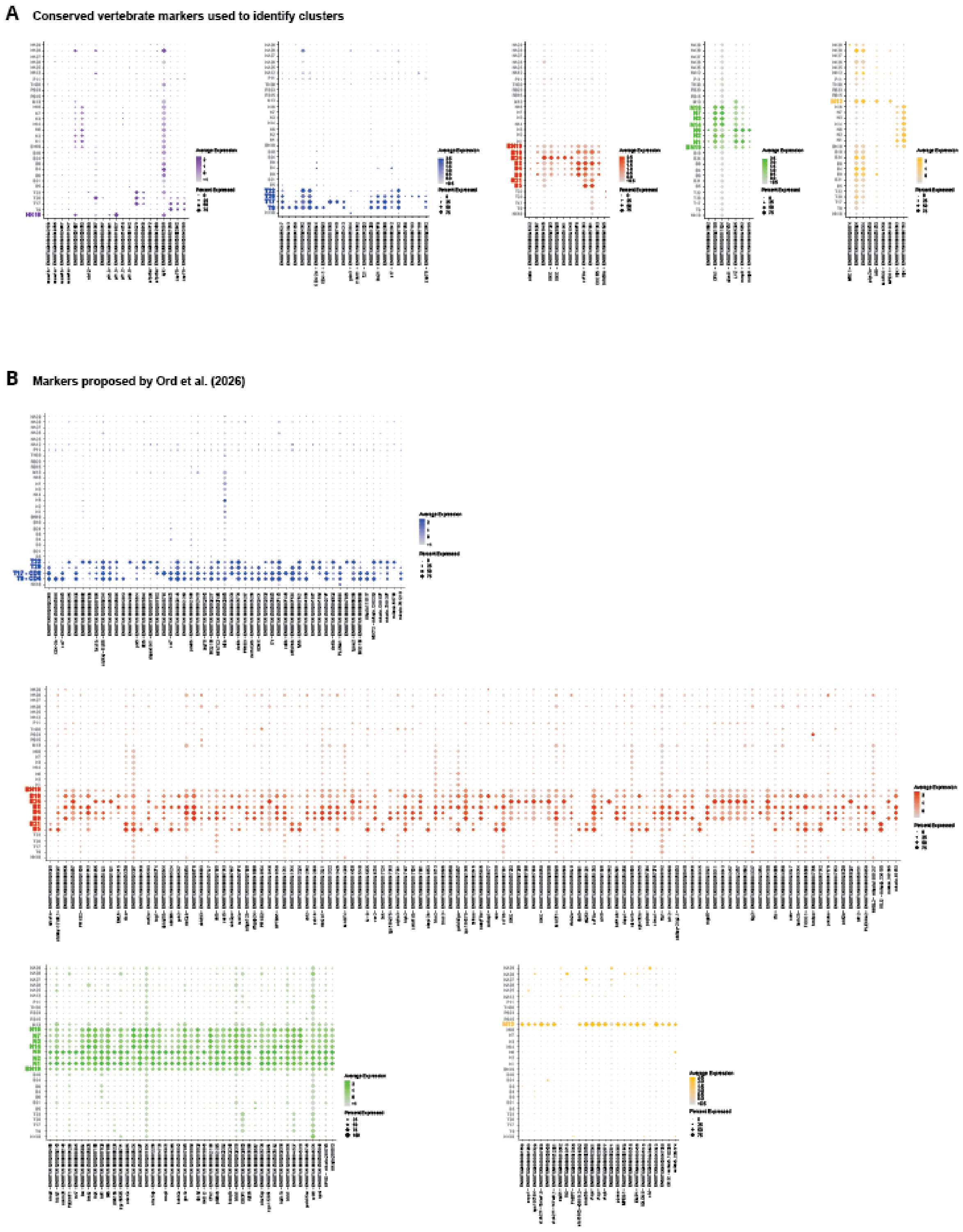


***Supplementary Figure 4. Expression of established and proposed marker genes across cell clusters.***

*(A) Expression of conserved vertebrate immune-cell markers used to support cluster annotation. (B) Expression of cell-type markers proposed by Ord et al. (2026). In both panels, dot size represents the percentage of cells within each cluster expressing the indicated gene, while colour intensity represents the average expression level among cells in that cluster.*

**Supplementary Figure 5.**

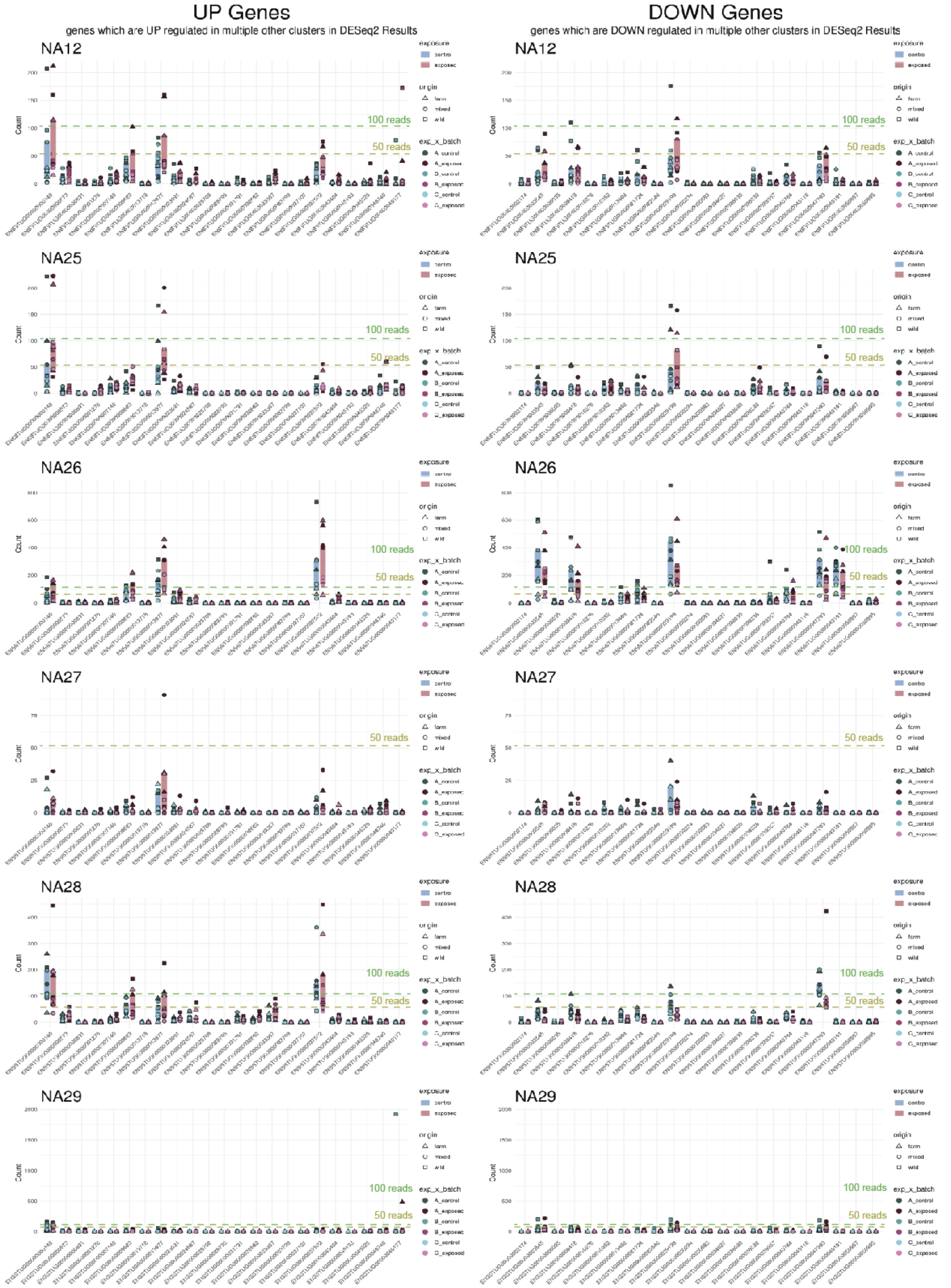


***Supplementary Figure 5. Expression of exposure-responsive genes in non-assigned cell clusters.***

*Raw read counts are shown for genes identified by DESeq2 as significantly upregulated (left column) or downregulated (right column) following exposure in annotated cell clusters. Olive and green dashed horizontal lines indicate read counts of 50 and 100, respectively, to facilitate comparison of expression levels across genes and clusters.*

**Supplementary Figure 6**

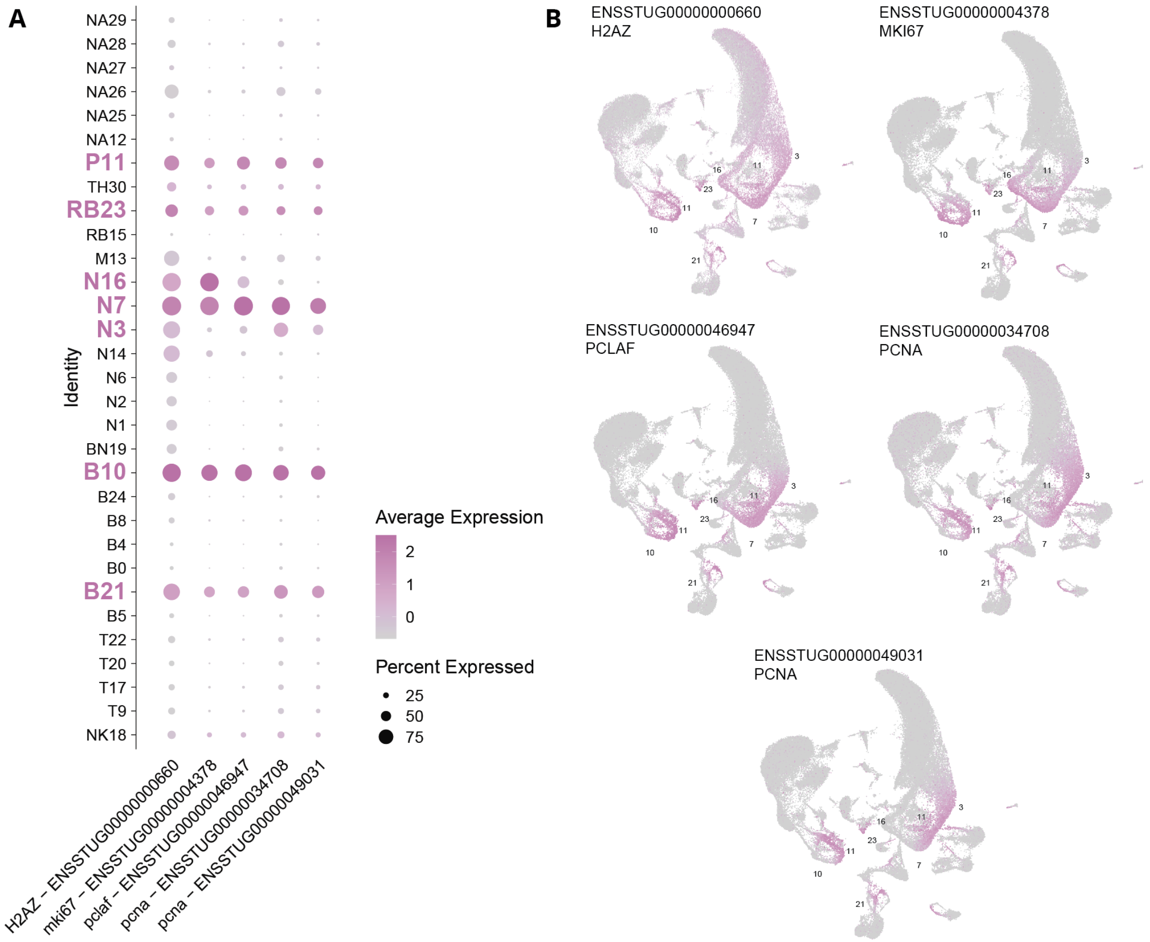


***Supplementary Figure 6. Expression of proliferation-associated genes across cell clusters.***

*(A) Dot plot showing the expression of the proliferation markers h2az, mki67, pclaf, and two pcna genes across all identified cell clusters. Dot size represents the percentage of cells within each cluster expressing the respective gene, while colour intensity indicates average expression. Cluster labels highlighted in magenta denote clusters exhibiting a proliferation-associated transcriptional signature.* ***(B)*** *UMAP projections showing the distribution of each proliferation marker across the complete dataset. Cells expressing the indicated gene are shown in purple, whereas other cells are shown in grey. Numbers indicate the identities of the proliferative clusters.*

**Supplementary Figure 7**

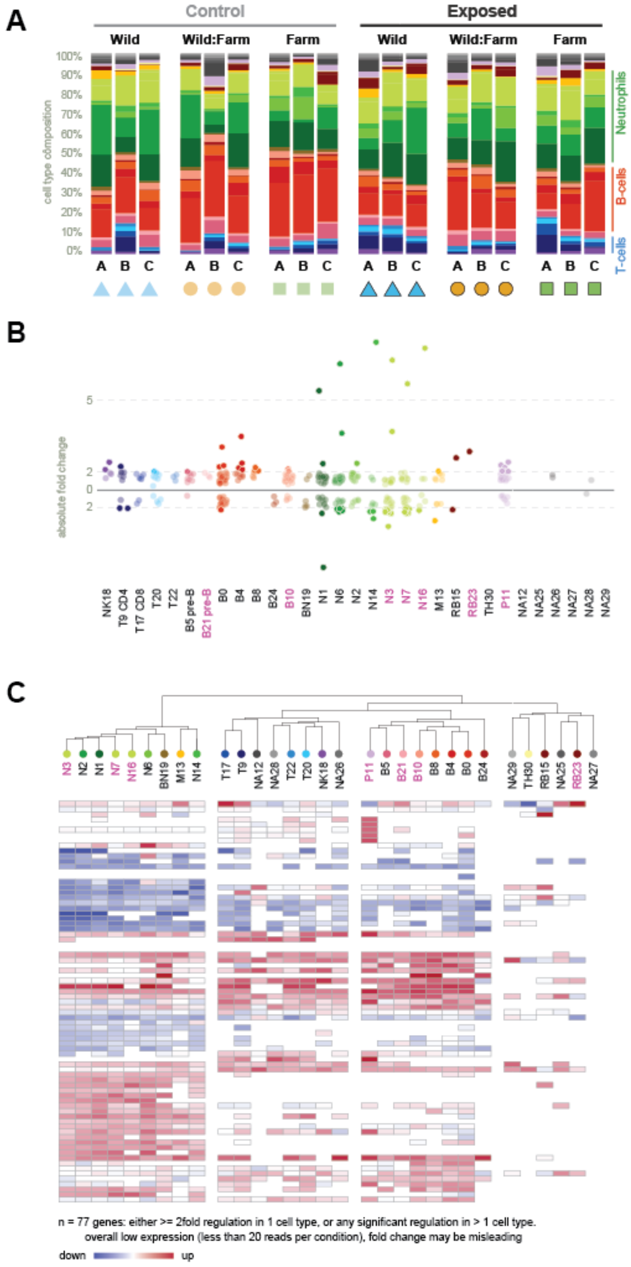


***Supplementary Figure 7. Effects of pathogen exposure on immune-cell composition and gene expression.***

*(A) Relative abundance of the identified cell clusters in control and exposed fish from the Wild, Wild:Farm, and Farm origins. Each stacked bar represents one individual. Colours indicate individual cell clusters, grouped into major immune-cell lineages. (B) Absolute fold changes in gene expression between exposed and control fish. Points represent individual genes. Horizontal reference lines indicate two- and five-fold changes. Cluster labels shown in magenta indicate cell types with a proliferation-associated transcriptional signature. (C) Heatmap showing exposure-associated gene regulation across cell clusters. Columns represent cell clusters and rows represent genes; clusters were hierarchically grouped according to their transcriptional responses. Red indicates increased and blue decreased expression in exposed relative to control fish, with colour intensity reflecting the magnitude of regulation. The heatmap includes 77 genes that showed either at least a two-fold change in one cell type or significant regulation in more than one cell type. For genes with fewer than 20 reads per condition, fold-change estimates should be interpreted cautiously.*

**Supplementary Figure 8**

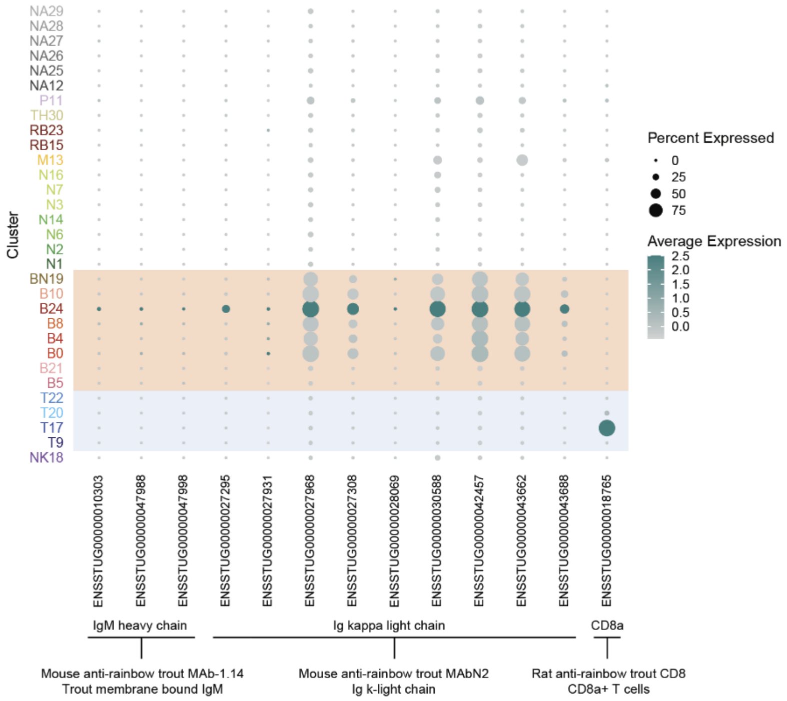


***Supplementary Figure 8. Expression of transcripts corresponding to antibodies used by Saura Martinez et al. (2026).***

*Dot size indicates the percentage of cells expressing each transcript, and colour intensity indicates average expression within each cluster. Candidate brown trout ighm genes were identified in Ensembl based on orthology to zebrafish ighm (ENSDARG00000096355). Neither candidate ighm transcript showed detectable expression in this dataset, for reasons that remain unclear. In contrast, immunoglobulin κ light-chain transcripts were expressed across several B-cell clusters, but with marked differences in prevalence and abundance, indicating that the corresponding B-cell antibody is unlikely to label all B-cell populations equally. cd8a expression was restricted to the expected T-cell cluster. Shaded areas indicate B-cell and T-cell populations.*
